# Systematic Epigenome Editing Captures the Context-dependent Instructive Function of Chromatin Modifications

**DOI:** 10.1101/2022.09.04.506519

**Authors:** Cristina Policarpi, Marzia Munafò, Stylianos Tsagkris, Valentina Carlini, Jamie A. Hackett

## Abstract

Chromatin modifications are linked with regulating patterns of gene expression, but their causal role and context-dependent impact on transcription remains unresolved. Here, we develop a modular epigenome editing platform that programmes nine key chromatin modifications – or combinations thereof – to precise loci in living cells. We couple this with single-cell readouts to systematically quantitate the magnitude and heterogeneity of transcriptional responses elicited by each specific chromatin modification. Amongst these, we show installing H3K4me3 at promoters causally instructs transcription activation by hierarchically remodeling the chromatin landscape. We further dissect how DNA sequence motifs influence the transcriptional impact of chromatin marks, identifying switch-like and attenuative effects within distinct *cis* contexts. Finally, we examine the interplay of combinatorial modifications, revealing co-targeted H3K27me3 and H2AK119ub maximise silencing penetrance across single-cells. Our precision perturbation strategy unveils the causal principles of how chromatin modification(s) influence transcription, and dissects how quantitative responses are calibrated by contextual interactions.

## INTRODUCTION

Understanding the molecular basis of gene regulation is a central challenge in modern biology. Regulation of eukaryotic transcription is guided by a complex interplay between transcription factors (TF), *cis* regulatory elements, and epigenetic mechanisms. The latter includes chromatin-based systems, and most prominently post-translational histone and DNA modifications. Such ‘chromatin modifications’ influence transcription activity via directly altering chromatin compaction, by acting as specific docking sites for ‘reader’ proteins, and/or by influencing transcription factor (TF) access to cognate motifs^1-3^. As a result, chromatin marks are thought to play a central regulatory role in deploying and propagating gene expression programs during development, whilst conversely, aberrant chromatin profiles are linked with gene mis-expression and pathology^4-6^.

The prominent role of chromatin modifications in genome regulation has spurred major initiatives to map their genome-wide distribution across healthy and disease cell types, revealing correlations with genomic features and transcription activity^7-11^. For example, histone H3 lysine 4 trimethylation (H3K4me3) is enriched at active gene promoters, H3K9me2/3 and H3K27me3/H2AK119ub are correlated with transcriptional repression, whilst active enhancers are co-marked by H3K4me1 and H3K27ac. However, whether the observed correlations indicate causation remains unclear. Indeed, depleting H3K4me1 or H3K27ac from embryonic stem cell (ESC) enhancers has only a relatively minor impact^12,13^. Moreover, the genomic landscape of activating histone modifications can be predicted and modulated by nascent transcription, implying marks such as H3K4me3 primarily reflect a consequence of gene expression^14,15^. To directly interrogate the functional relevance of epigenetic marks, perturbation strategies have been widely deployed, often by manipulating chromatin-modifying enzymes or histone residues ^5,16,17^. However, whilst insightful, such global approaches affect the entire (epi)genome simultaneously, and thus render it challenging to distinguish *direct* from *indirect* effects. Indeed, chromatin-modifying enzymes also have multiple non-histone substrates ^18,19^ and non-catalytic roles ^20,21^, whilst residues typically acquire multiple modifications, which all complicates interpretation of their loss-of-function. Thus, the extent to which chromatin modifications *per se* causally instruct gene expression states remains unresolved.

A deeper understanding of the functional role of epigenetic modifications on DNA-templated processes would be facilitated by development of tools for precision chromatin perturbations. Epigenome editing technologies that enable manipulation of specific chromatin states at target loci have recently emerged, primarily based around programmable dCas9-fusion systems ^22,23^. For example, P300 and HDAC3 have been fused to dCas9 to deposit or remove histone acetylation ^24,25^. Further approaches have engineered dCas9 systems that specifically edit DNA methylation, H3K27me3, H3K27ac, H3K4me3, and H3K79me3 ^26-34^. Such pioneering studies have revealed proof-of-principle that altering the epigenome can be sufficient to induce at least some changes in gene expression. However, the transcriptional responses to specific marks are generally modest, if at all, and register at only a restricted set of target genes. This may partly reflect technical limitations in depositing physiological levels of chromatin marks, but likely also implies their functional impact varies depending on context-dependent influences. Indeed, there is increasing appreciation that factors such as underlying DNA motifs/variants and the cell type-specific repertoire of TF will all modulate the precise impact of a chromatin modification at a given locus ^35,36^. Thus, beyond the principle of causality, it is important to deconvolve the degree to which specific chromatin marks affect transcription levels *quantitatively* (as opposed to an ON/OFF toggle), how DNA sequence context influences this, and the hierarchical relationships involved.

Here, we develop a suite of modular epigenetic editing tools to systematically programme nine biologically-important chromatin modifications to specific loci at physiological levels. By coupling this with single-cell readouts, we capture the causal and quantitative impact of each modification on transcription. We further show that epigenetic marks are linked to each other by specific hierarchical interplays, and function combinatorially to promote robustness in transcriptional responses. We finally dissect how the impact of chromatin marks is influenced by sequence motifs and TF binding, identifying switch-like functionality within different *cis* contexts. The output is a framework for quantifying the instructive role of chromatin modifications, and their functional interplay with other regulatory mechanisms.

## RESULTS

### A toolkit for precision programming of chromatin modifications at endogenous loci

We sought to engineer a modular epigenetic editing system that can programme *de novo* chromatin modification(s) to specific target loci at physiological levels. To achieve this, we exploited a catalytically <u>d</u>ead *Cas9* (dCas9) fused with a tail-array of five GCN4 motifs (dCas9^GCN4^) ^37,38^. This tethers up to *five* scFV-tagged epigenetic ‘effectors’ to genomic targets, which amplifies editing activity (Fig 1A). To programme a broad range of specific chromatin modifications, we built a library of effectors each comprising the <u>c</u>atalytic <u>d</u>omain (CD) of a DNA-or histone-modifying enzyme linked with scFV (collectively: CD^scFv^). By isolating the catalytic domain, we can exclude confounding effects of tethering entire chromatin-modifying proteins, which can exert non-catalytic regulatory activities. The toolkit includes catalytic cores that deposit H3K4me3 (*Prdm9*-CD^scFv^), H3K27ac (*p300*-CD^scFv^), H3K79me2 (*Dot1l*-CD^scFv^), H3K9me2/3 (*G9a*-CD^scFv^), H3K36me3 (*Setd2*-CD^scFv^), DNA methylation (*Dnmt3a3l*-CD^scFv^), H2AK119ub (*Ring1b*-CD^scFv^) and full-length enzymes that write H3K27me3 (*Ezh2-*FL^scFv^) and H4K20me3 (*Kmt5c*-FL^scFv^) (Fig 1A). As further controls, we generated catalytic point-mutants for each CD^scFV^ effector (mut-CD^scFv^), which specifically abrogates their enzymatic activity (Fig S1A). Our strategy therefore enables direct assessment of the functional role of the deposited chromatin mark *per se*.

**Figure 1.**
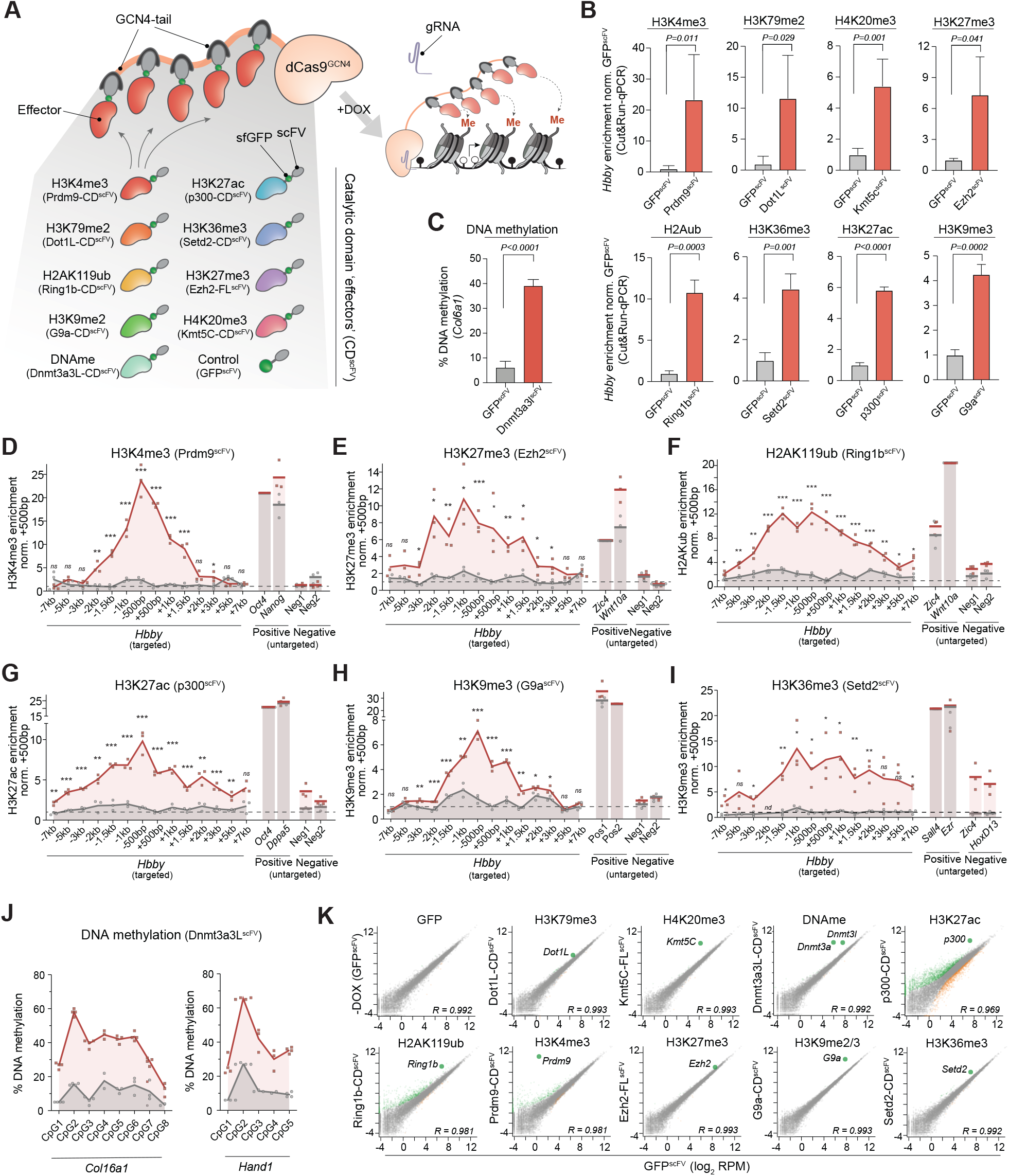
A modular toolkit for precisely programming chromatin states. **(A)** Schematic of the modular epigenetic editing platform. Upon DOX-induction, dCas9^GCN4^ recruits five copies of chromatin-modifying effector(s) or control GFP^scFV^ to target loci via a specific gRNA. **(B)** Relative abundance of the indicated histone modification at *Hbby* assayed by either CUT&RUN-or ChiP-qPCR (H3K36me3, H3K79me2), following epigenetic-editing or control GFP^scFV^ recruitment in ESC for seven days. Shown is the mean of three biological replicates; error bars indicate S.D. **(C)** Histogram showing mean DNA methylation installed at the unmethylated *Col16a1* promoter by epigenetic editing, determined by bisulfite pyrosequencing in triplicate biological samples. **(D-I)** Relative abundance of the indicated histone modification across the *Hbby* locus after epigenetic programming with a specific CD^scFV^ (red line) or control GFP^scFV^ (grey line), assayed by CUT&RUN-qPCR. Mean enrichment across a ∼14kb region centered on gRNA binding sites is shown for biological triplicate editing, as well for endogenous positive and negative loci for each mark. **(J)** Percentage DNA methylation at CpG dinucleotides across the *Col16a1* and *Hand1* promoters in triplicate experiments. **(K)** Scatter plots showing limited global gene expression changes following 7 days targeted deposition of the indicated epigenetic mark at the *Hbby* locus, relative to control GFP^scFV^ targeting. Differentially expressed genes are indicated in green/or-ange. Grey dots indicate unaffected genes. *P-values* in all panels are calculated by unpaired t-test. *\*P<0*.*05 **P<0*.*01, ***P<0*.*001*.

We engineered the system to be doxycycline (DOX)-inducible to facilitate dynamic ON-OFF epigenetic editing. Moreover, all CD^scFv^ effectors are tagged with superfolder GFP (sfGFP) to monitor protein stability, to track dynamics, and to isolate epigenetically edited populations (Fig S1B). Locus-specific editing is directed by an enhanced gRNA scaffold (AT-flip, extended stem loop) with tagBFP ^39^. Finally, up to three nuclear localisation sequences (NLS) were incorporated into each effector, since we found two NLS were routinely insufficient for robust nuclear accumulation, for example for *Dot1l*-CD^scFv^ (Fig S1C).

To test the capacity to programme specific *de novo* epigenetic states, we introduced dCas9^GCN4^ and each CD^scFv^ into mouse ESC *via* piggyBac, and targeted the endogenous *Hbby* locus. Following induction with DOX, we observed that each effector directed highly significant deposition of its cognate histone modification relative to recruitment of GFP^scFv^ alone, using quantitative CUT&RUN-qPCR. This includes *de novo* establishment of H3K27ac (*P<0*.*0001*), H3K4me3 (*P=0*.*011*), H3K79me2 (*P=0*.*029*), H4K20me3 (*P=0*.*001*), H3K27me3 (*P=0*.*041*), H2AK119ub (*P=0*.*0003*), H3K36me3 (*P=0*.*001*), H3K9me2/3 (*P=0*.*0002*) (Fig 1B). Comparable chromatin mark targeting was independently achieved using either one or three gRNAs together (Fig S1D). We also found highly significant programming of DNA methylation (*P<0*.*0001*) upon recruitment of *Dnmt3a3L*-CD^scFv^ (Fig 1C).

To determine the quantitative level (amplitude) and spreading (domain breadth) of induced epigenetic editing, we assessed enrichment across the entire *Hbby* locus. We typically observed a peak of each programmed histone modification centered on the gRNA binding sites, with significantly modified domains extending more than 2kb either side, which likely reflects the flexible tail-array structure of dCas9^GCN4^. Enrichment of targeted histone modifications ranged from 7 to >20-fold over background (Fig 1D-I) and importantly, in most cases were of comparable quantitative levels to strong positive peaks within the genome. For example, programmed H3K4me3 enrichment (*Prdm9*-CD^GFP-scFv^) at *Hbby* was equivalent to highly-marked *Oct4* and *Nanog* promoters (Fig 1D), whilst polycomb marks H3K27me3 (*Ezh2*-FL^scFv^) and H2AK119ub (*Ring1b*-CD^scFv^) were *de novo* deposited with similar enrichments as endogenous polycomb targets *Zic4* and *Wnt10a* (Fig 1E-F). Moreover, *de novo* H3K36me3, H3K79me3, and H4K20me3 were comparable with endogenous peaks, whilst H3K9me2/3 and H3K27ac were deposited at levels moderately lower than control targets (Fig 1G-I & S1E). Finally, up to 60% DNA methylation was installed at previously unmethylated promoters (Fig 1J). Taken together, our inducible epigenetic editing toolkit programmes specific chromatin modifications to target loci at physiologically-relevant levels, in both amplitude and domain breadth.

We did not detect OFF-target chromatin deposition at negative (non-targeted) genomic loci with most effectors (Fig 1D-I & S1E), implying the strategy facilitates specific ON-target chromatin mark editing. To confirm this further, we performed RNA-seq following DOX-induction of each CD^scFV^. We observed only minor changes in global gene expression following activation, with the top hit invariably mapping to the endogenous domain of the activated CD^scFV^ chromatin-modifier (Fig 1K & S2A). An exception is p300-CD^scFV^, for which we observed a global expression impact. To mitigate this going forward we limited p300-CD^scFV^ induction levels by using 20-fold lower DOX. Overall, the data suggest intrinsic OFF-target activity and/or indirect effects is minimised with our modular CD^scFV^ recruitment design. Thus, we have developed a flexible epigenetic editing toolkit capable of programming high levels of nine biologically important chromatin modifications to specific endogenous loci. The system includes multiple controls to isolate the effects of chromatin modifications *per se*, is compatible with combinatorial mark targeting, and can track temporally-resolved responses and epigenetic memory. This collectively enables a systematic analysis of the causal function of distinct chromatin states through precision perturbations, without confounding global effects.

### Chromatin modifications instruct transcriptional outputs at single-cell resolution

To investigate the direct regulatory role of chromatin modifications on transcriptional control, we initially engineered a reporter system, which facilitates quantitative single cell readouts. Here we embedded the endogenous *Ef1a* core promoter (212bp) into a contextual DNA sequence (∼3kb) selected from the human genome to be feature-neutral on the basis of the following criteria: it carries no transposable elements, is 50% GC, and has minimal transcription factor (TF) motifs (Fig 2A). This design enables the impact of introducing specific genetic motifs to be tested in future (*see* Fig 4). We inserted this ‘reference’ (REF) reporter into two distinct genomic locations, chosen to be either permissive (Chr9) or non-permissive (Chr13) for transcriptional activity (Fig 2A). Consistently, knock-in to the permissive locus supported strong expression (ON), whereas the non-permissive landing site resulted in minimal activity (OFF), which partially reflects acquisition of polycomb silencing (Fig 2B & S2B). These identical reporters residing within distinct genomic locations thus enable controlled assessment of both activating and repressive activity of an induced chromatin modification on the same underlying DNA sequence.

**Figure 2.**
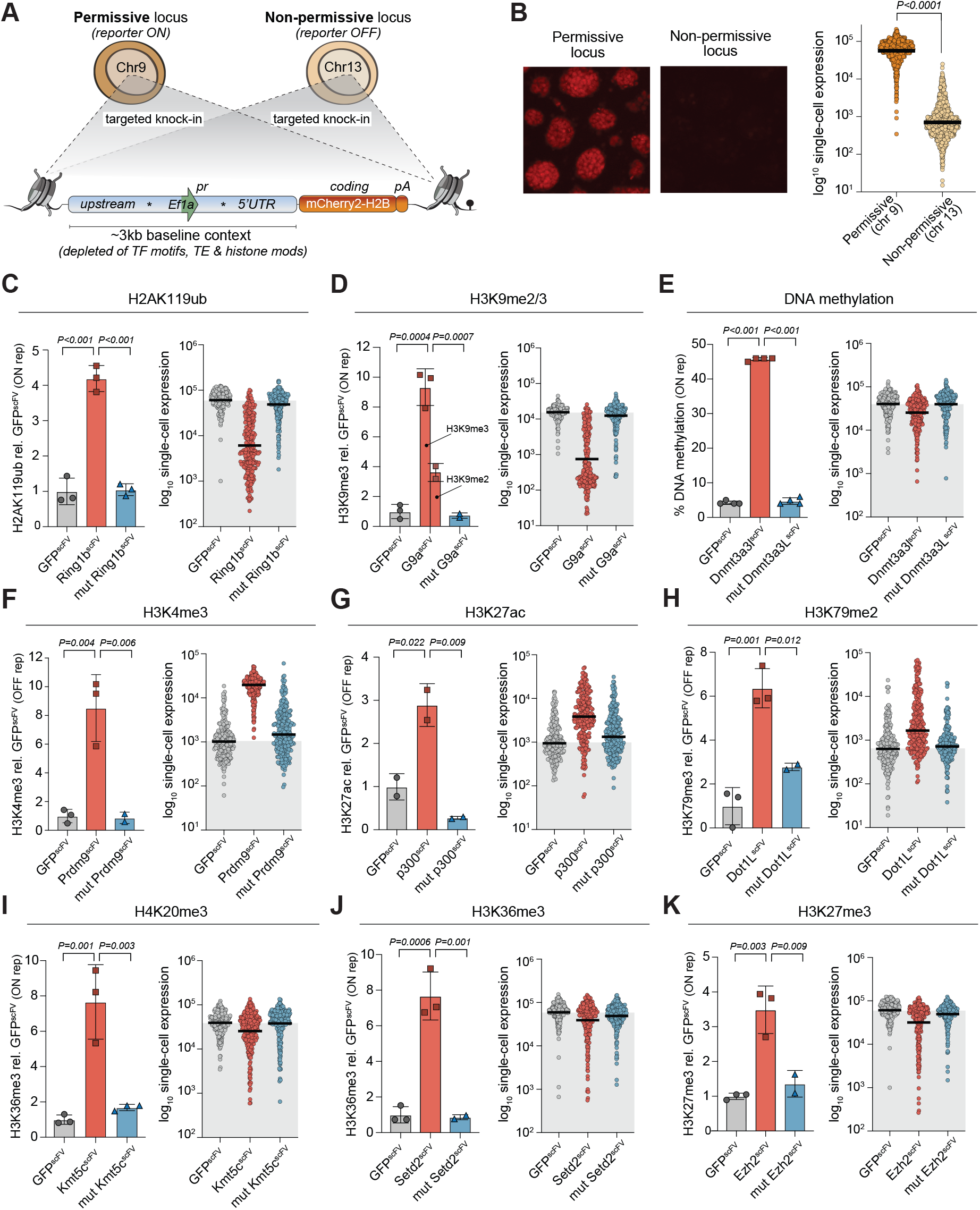
Distinct chromatin modifications causally instruct transcriptional responses. **(A)** Schematic depicting the structure of the Reference reporter and its targeted integration into either a transcriptionally permissive (chr9, ON) or non-permissive (chr13, OFF) locus. Stars indicate gRNA target sites within the neutral DNA context. (**B)** Representative fluorescence images (left) and dot plot (right) from quantitative flow cytometry showing activity of the Reference reporter when integrated into either the permissive or non-permissive locus. Bars denote the geometric mean. *P-value* is by unpaired t-test. **(C-K)** Programming of a specific chromatin modification (left) and transcriptional responses in single-cells (right) for **(C)** H2AK119ub, **(D)** H3K9me2/3, **(E)** DNA methylation, **(F)** H3K4me3 **(G)** H3K27ac **(H)** H3K79me2 **(I)** H4K20me3 **(J)** H3K36me3 **(K)** H3K27me3. Left in each panel: Histogram showing the relative enrichment of the indicated chromatin modification after targeting control GFP^scFV^ (grey bar), wild-type CD^scFV^ (red bar) or catalytic-inactive mut-CD^scFV^ (blue bar) for seven days. Displayed is the mean of at least two independent quantitations by CUT&RUN-or ChIP-qPCR. Error bars represent S.D. Right: Dot plot showing log_10_ expression (mCherry2) in response to epigenetic editing of the indicated chromatin mark. Each data-point represents a single cell expression value, bars denote the geometric mean in the population. *P-values* are calculated by one-way ANOVA with Tukey’s multiple test correction. *\*P<0*.*05 **P<0*.*01, ***P<0*.*001*.

We targeted each CD^scFV^ to each reporter, and initially confirmed highly significant programming of the expected chromatin modification relative to control GFP^scFV^ (Fig 2C-K, left panels). Importantly, targeting catalytic-mutant effectors (mut-CD^scFV^) did not change the chromatin state (Fig 2C-K). We therefore moved to assess the functional impact of each programmed mark on transcription quantitatively and in single cells via flow cytometry. Using this sensitive strategy, we were able to detect that deposition of each tested chromatin modification has the potential to instigate at least some quantitative transcriptional response. Based on this, we grouped chromatin marks into three functional categories; (i) Modifications that instruct transcriptional *repression*, with penetrance across the majority fraction of cells; (ii) Modifications that trigger transcription *activation*, with high penetrance; (iii) Modifications that have *subtle* and/or *partially penetrant* transcriptional effects.

The first group is characterised by the polycomb repressive complex 1 (PRC1) modification H2AK119ub, and the heterochromatin mark H3K9me2, which is endogenously converted to H3K9me3. We find that *de novo* deposition of either H2AK119ub or H3K9me2/3 is sufficient to drive transcriptional silencing of the permissive (ON) reporter >100-fold in some cells, with average repression across the population exceeding 10-fold (geometric mean) (Fig 2C-D, right panels). Moreover, whilst there was heterogeneity, >98% of cells shifted expression below the average level of control GFP^scFV^ in response to either H2AK119ub or H3K9me2/3. DNA methylation is also included here as its deposition resulted in a modest but penetrant population shift in expression, averaging 1.9-fold (±0.1 S.D) repression (Fig 2E & S3A). Of note, targeting mut-Ring1b-CD^scFV^, mut-G9a-CD^scFV^ or mut-Dnmt3a3l-CD^scFV^ had no significant impact on expression (Fig 2C-E). This indicates that the H2AK119ub and H3K9me2/3 marks *per se* are sufficient to causally instruct robust silencing of an active promoter, whilst partial (∼50%) DNA methylation causes moderate albeit detectable repression. Moreover, deposition of H3K9me2/3 or DNA methylation at the non-permissive (OFF) reporter was capable of inducing even further silencing, potentially via synergising with polycomb (Fig S3A).

The second response group comprised chromatin modifications sufficient to induce quantitative transcriptional activation, when deposited at a repressed promoter. These were represented by H3K4me3, H3K27ac and H3K79me2. Programming each mark triggered a reproducible population shift leading to 18.1-fold (±3.8), 3.5-fold (±0.2), and 2.4-fold (±0.4) increased expression, respectively, with some cells within the population activating >50-fold over GFP^scFV^ (Fig 2F-H). Targeting catalytic inactive mut-Prdm9-CD^scFV^, mut-p300-CD^scFV^, or mut-Dot1l-CD^scFV^ did not elicit transcriptional responses. Neither H3K79me2 nor H3K27ac deposition at the active (ON) locus further enhanced its expression, whereas additional H3K4me3 shifted cells into a narrow band of maximal expression (Fig S3B). These data indicate acquisition of promoter H3K27ac, and to a lesser extent H3K79me2, can promote transcriptional activation of a repressed locus, albeit relatively modestly for the latter (Fig 2G-H). Furthermore, these data surprisingly imply H3K4me3 *per se* has the capacity to instigate strong transcription upregulation (Fig 2F).

The third functional group of chromatin modifications elicited regulatory impacts that led to variable or weak repressive responses at the active locus. Amongst these, targeted deposition of H4K20me3, H3K36me3 and H3K27me3 instigated a degree transcriptional repression at the population level. This amounted to 1.6-fold (±0.3), 1.2-fold (±0.1), and 1.5-fold (±0.1) (geometric mean), respectively, with the relevant catalytic mutant CD^scFV^ controls bearing no effect (Fig 2I-K). Notably however, H3K27me3, H3K36me3 and H4K20me3 were distinguished by the imposition of strong silencing in a highly heterogeneous manner (>50-fold in some cells) but with the majority of cells remaining within the original expression level, resulting in a broad distribution of transcriptional responses (Fig 2I-K & S3C). Because other equivalently-enriched modifications elicited more penetrant impacts, these heterogeneous responses likely reflect biological rather than technical outcomes, such as dynamic competition between *de novo* marks and the transcription machinery. Irrespective, these data support the principle that the acquisition of H4K20me3, H3K36me3 or H3K27me3 marks are capable of impacting transcriptional responses heterogeneously within a cellular population, albeit subtly.

Finally, we assessed how programming each modification affects chromatin accessibility. In all cases, we found promoter accessibility is well correlated with the directionality of gene expression induced by epigenetic editing, further supporting the impact of modifications on transcription (Fig S4A). Indeed, we also observed a dose-dependent correlation between the induction level of the epigenetic editing machinery and transcriptional responses, suggesting gene activity can be tuned with chromatin modifications (Fig S4B-E). In summary, by exploiting a sensitive single-cell readout and precision epigenome editing, we capture that *de novo* epigenetic marks can causally instigate quantitative changes in gene expression. We report the magnitude and nature of these changes, which vary from robust, to subtle and/or heterogenous, to non-functional, depending on the identity of the mark and the genomic context. These data thus support the principle that each of the nine biologically-relevant chromatin modifications tested here has the *potential* to directly influence transcription output, when measured at an appropriate quantitative and single-cell resolution.

### H3K4me3 can direct transcription initiation

Amongst the most striking impacts of precision epigenetic editing was that of H3K4me3 deposition, which induced robust reporter activation (Fig 2F). H3K4me3 is universally correlated with transcriptional activation, yet whether it is responsible for instructing expression or merely a consequential marker is intensely debated ^14,40^. Indeed, current paradigms suggest H3K4me3 contributes to preventing gene silencing, but does not in itself instigate gene activation ^41^. To probe the functional impact of H3K4me3 further, we generated ESC carrying homozygous knock-in Y2602A point catalytic mutations (CM) in the key H3K4 methylase *Mll2*, to specifically disrupt its enzymatic activity (*Mll2*^*CM/CM*^). This enables the loss-of-function of the H3K4me3 mark *per se* to be assessed without confounding issues associated with deletion of MLL2 protein/complexes. CUT&RUN identified a cluster of 3,332 MLL2-dependent H3K4me3 promoter peaks that are lost in *Mll2*^*CM/CM*^ ESC, whilst 13,477 promoters retain H3K4me3, likely due to redundant H3K4me3 modifiers (Fig 3A). Amongst genes that lose H3K4me3, expression of 458 (90%) is significantly downregulated (*P(adj)<0*.*05*), with only 53 (10%) upregulated, consistent with H3K4me3 *per se* playing a role in preventing gene repression. Indeed, promoter clusters with no H3K4me3 change are equally likely to be up-or down-regulated (Fig 3A & S5A-B). Moreover, promoters that lose H3K4me3 in *Mll2*^*CM/CM*^ ESC also become depleted of H3K27ac and exhibit a gain of diffuse H3K27me3 domains (Fig S5C-D). This implies that specific removal of H3K4me3, but not MLL2, unmasks the potential for silencing of a subset of genes that were previously active.

**Figure 3.**
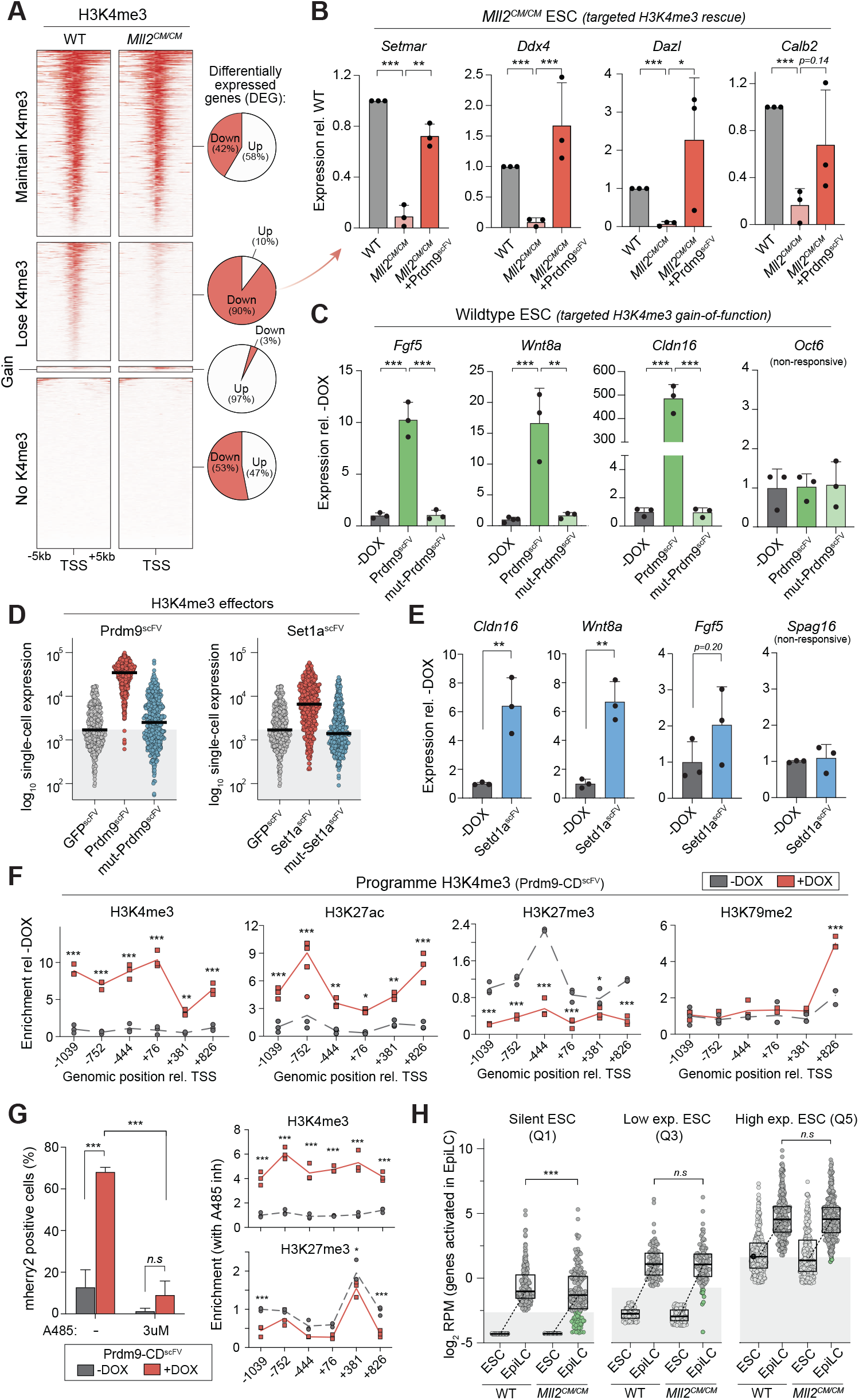
*De novo* H3K4me3 triggers transcription upregulation. **(A)** H3K4me3 enrichment over the transcrip-tional start site (TSS) ±5kb in wild-type and *Mll2*^*CM/CM*^ ESC, stratified according to H3K4me3 change in *Mll2*^*CM/CM*^ ESC. The percentage of up-or down-differentially expressed genes in each category is shown. **(B)** Bar plots showing expression of indicated genes in wild-type, *Mll2*^*CM/CM*^ and *Mll2*^*CM/CM*^ +Prdm9^scFV^ ESC, in which H3K4me3 has been programmed back to a repressed endogenous promoter. Shown is mean of three biological replicates by qRT-PCR. Error bars represent S.D and significance of rescue is calculated by unpaired t-test. **(C)** Bar plots of endogenous gene expresion in wild-type ESC or upon targeting with Prdm9^scFV^ to programme H3K4me3, or mut-Prdm9^scFV^. Data is the mean of biological triplicates; error bars represent S.D. Significance is calculated by one-way ANOVA with Tukey’s correction. (**D)** Dot plots showing expression at the OFF reporter after targeting with distinct H3K4me3 effectors Prdm9^scFV^ (left) or Set1a^scFV^ (right). Each data point is a cell and bars denote the geometric mean. (**E)** Bar plots of gene expression in wild-type ESC targeted with Set1a^scFV^ or untargeted (-DOX), by RT-qPCR from biological triplicates. Error bars S.D with significance by unpaired t-test. **(F)** Epigenetic landscape response at the OFF reporter before (-DOX) and after (+DOX) specific H3K4me3 programming. Indicated indicated histone modification enrichment across ∼2kb in triplicate technical samples, with significance calculated by unpaired t-test. **(G)** Left: Bar plots showing the percentage of mCherry positive cells is restricted after (+DOX) H3K4me3 installation by Prdm9^scFV^ in the presence of p300 inhibitor A485. Data is biological triplicate, error bars represent S.D. *P-values* calculated by two-way ANOVA. Right: Relative abundance of the indicated histone modifications after programming H3K4me3 (+DOX) in presence of A485. **(H)** Dot plots showing log expression of genes (grey dots) that are normally activated in EpiLC stratified into quintiles (Q) according to their initial expression level in ESC. Box plots indicate median and interquartile range. Genes that fail to activate in *Mll2*^*CM/CM*^ EpiLC are shown in green. Significance is calculated by unpaired t-test, *\*P<0*.*05 **P<0*.*01, ***P<0*.*001*.

To distinguish whether H3K4me3 simply safeguards against silencing versus whether H3K4me3 is capable of *instigating* transcriptional upregulation, we next programmed H3K4me3 back to eight genes that became repressed in *Mll2*^*CM/CM*^ cells, of which seven lose H3K4me3. Upon DOX-induction of Prdm9-CD^scFV^ to restore H3K4me3, we found all seven genes showed a trend of reactivation, with 5 of 7 reaching significant transcriptional rescue, including *Setmar, Dazl* and *Ddx4* (Fig 3B & S6A). In contrast, the control *Pldn* gene, which is downregulated without H3K4me3 loss in *Mll2*^*CM/CM*^ cells, exhibited no reactivation (Fig S6A). This suggests that acquisition of H3K4me3 at promoters can activate endogenous genes that were previously expressed, prior to genetically-induced depletion of H3K4me3.

To examine whether H3K4me3 can also instigate expression of genes that are never active in a given cell type, we targeted H3K4me3 to eight randomly selected silent promoters in naïve ESC. Installation of H3K4me3 resulted in significant activation at 3 out of 8 of these genes, with maximal upregulation reaching >400-fold at *Cldn16* (Fig 3C). Importantly, targeting the catalytically inactive mut-Prdm9CD^scFV^ had no detectable impact. These data support the conclusion that forced programming of H3K4me3 at promoters can overcome silencing to induce *de novo* transcription - at least at some genes - and that this reflects the activity of the H3K4me3 mark itself.

To further examine whether H3K4me3 *per se* rather than the Prdm9-CD^scFV^ effector can instruct transcription, we generated an independent H3K4me3 effector based on the catalytic core of *Set1a* (Set1A-CD^scFV^). We used our modular dCas9^GCN4^ system to target compound Set1A-CD^scFV^ to the OFF reporter, which triggered robust activation amongst a significant fraction of cells (Fig 3D). Indeed, >85% cells express above control average in response to H3K4me3, with 3.3-fold (±0.3 S.D) increased transcription across the population. The catalytic-inactive mut-Set1A-CD^scFV^effector had no impact (Fig 3D). Of note, the gradated level of activation induced by each effecter (Prdm9-CD^scFV^ > Set1A-CD^scFV^) correlated with the amount of H3K4me3 they each deposited (Fig S6B), suggesting a dosedependent impact of H3K4me3. Indeed, analysis of cells that failed to activate revealed they still acquire H3K4me3, but that responsive cells acquire more H3K4me3 (Fig S6C), implying a threshold level triggers a switch into active transcription at the single-cell level. Finally, we also targeted endogenous genes with Set1A-CD^scFV^, and again observed a significant transcriptional activation of some (2/4) upon H3K4me3 deposition (Fig 3E). Taken together, independent targeted gain-of-function approaches support the principle that sufficient H3K4me3 can activate productive transcription at otherwise silent promoters. Furthermore, the data show that in some instances, *de novo* H3K4me3 is not sufficient to activate transcription.

### Developmental role of H3K4me3

We next investigated the potential mechanisms through which H3K4me3 operates, by initially asking whether *de novo* H3K4me3 directs remodelling of the local chromatin landscape. We found that programming H3K4me3 to the OFF reporter caused a highly significant secondary recruitment of H3K27ac (Fig 3F). In parallel H3K27me3 is evicted by H3K4me3 deposition and there is a gain in promoter accessibility, whilst H3K79me2 remains largely unaltered (Fig 3F). This suggests *de novo* H3K4me3 establishment directly influences the balance of distinct H3K27 modifications, and more generally facilitates promoter acetylation. Because histone acetylation is linked with active transcription, we asked whether its recruitment is necessary for H3K4me3-mediated effects. We programmed H3K4me3 to the OFF reporter with or without the specific p300/CBP inhibitor, A485, which specifically blocks its acetyltransferase activity, including against H3K27ac, H3K18ac and H2B ^42^. A485 did not affect efficient programming of H3K4me3 (Fig 3G). However, the presence of A485 (3uM) restricted subsequent H3K4me3-mediated activation to <10% of cells, compared to ∼70% in no-inhibitor controls (Fig 3G & S6D). Programming H3K4me3 in the presence of A485 also largely blocked displacement of H3K27me3. This supports a hierarchical model whereby *de novo* H3K4me3 may functionally operate, at least partially, via directly or indirectly facilitating promoter acetylation and evicting epigenetic silencing systems.

To examine the potential physiological role for H3K4me3 in contributing to gene activation programmes during development, we induced differentiation of naive *Mll2*^*CM/CM*^ ESC into formative epiblast-like cells (EpiLC). This triggers 1,380 genes to undergo robust upregulation (*p(adj)<0*.*05; log*_*2*_*(FC)>2*) in wildtype cells. The majority of these genes activated normally in *Mll2*^*CM/CM*^ EpiLC, and indeed naïve and formative markers exhibited appropriate changes, suggesting MLL2-mediated H3K4me3 is not requisite for EpiLC cell fate transition (Fig S7A-C). However, by stratifying upregulated EpiLC genes into quintiles based on their initial expression level in ESC, we found that H3K4me3 *per se* appears necessary for activation of a subset of genes that are silent in ESC and then induced *de novo* in EpiLC (Fig 3H & S7D). Specifically, genes in the lowest quintile (Q1) of ESC expression fail to activate as expected in *Mll2*^*CM/CM*^ EpiLC (*P=0*.*0028*), whereas those genes that are already weakly or fully expressed in ESC (Q3-Q5), are competent to be upregulated (Fig 3H). For example, *Col1a2, Spon1* and *Lrp1b* normally acquire H3K4me3 and lose H3K27me3 coincident with upregulation in EpiLC, but fail to be appropriately activated in *Mll2*^*CM/CM*^ EpiLC that lack H3K4me3 catalytic activity (Fig S7D-E). This data collectively implies H3K4me3 contributes to initiating transcriptional activation of a subset of genes during cell fate transition.

In summary, our precision epigenetic editing strategy demonstrates that *de novo* H3K4me3 installation is sufficient to remodel the chromatin landscape and instigate transcriptional upregulation, at least at some genes, rather than simply reflecting a consequence of activity.

### Epigenetic marks interact with *cis* genetic motifs to modulate transcription

The precise functional impact of a given histone modification is likely dependent on contextual interactions, including with the underlying DNA sequence features at each promoter. We therefore next used our epigenetic editing strategy to investigate the interplay between DNA sequence variants and chromatin function. We generated a repertoire of reporters wherein each comprises the identical ∼3kb baseline sequence derived from the reference (REF) reporter (Fig 2A), but is distinguished by insertion of several short DNA motifs (8-14bp), thus establishing an allelic series. We selected motifs corresponding to binding sites of specific transcription factors (OCT4, OTX, MYC, GATA) or that impact chromatin architecture either indirectly through the recruitment of architectural proteins (CTCF, YY1) or directly via formation of G-quadruplexes (G4-U, G4-D) ^43,44^ (Fig 4A & Fig S8A). We knocked-in each reporter, which only differ by a few base pairs, into both the permissive (ON) and nonpermissive (OFF) genomic landing sites (*see* Fig 2A). Most motifs did not impact baseline expression levels, albeit inclusion of CTCF, G4-U or YY1 motifs decreased expression at the permissive locus, partly due to increased heterogeneity (Fig 4B). Overall, we generated a series of reporters that carry specific DNA sequence variants, within highly-controlled genomic environment(s).

**Figure 4.**
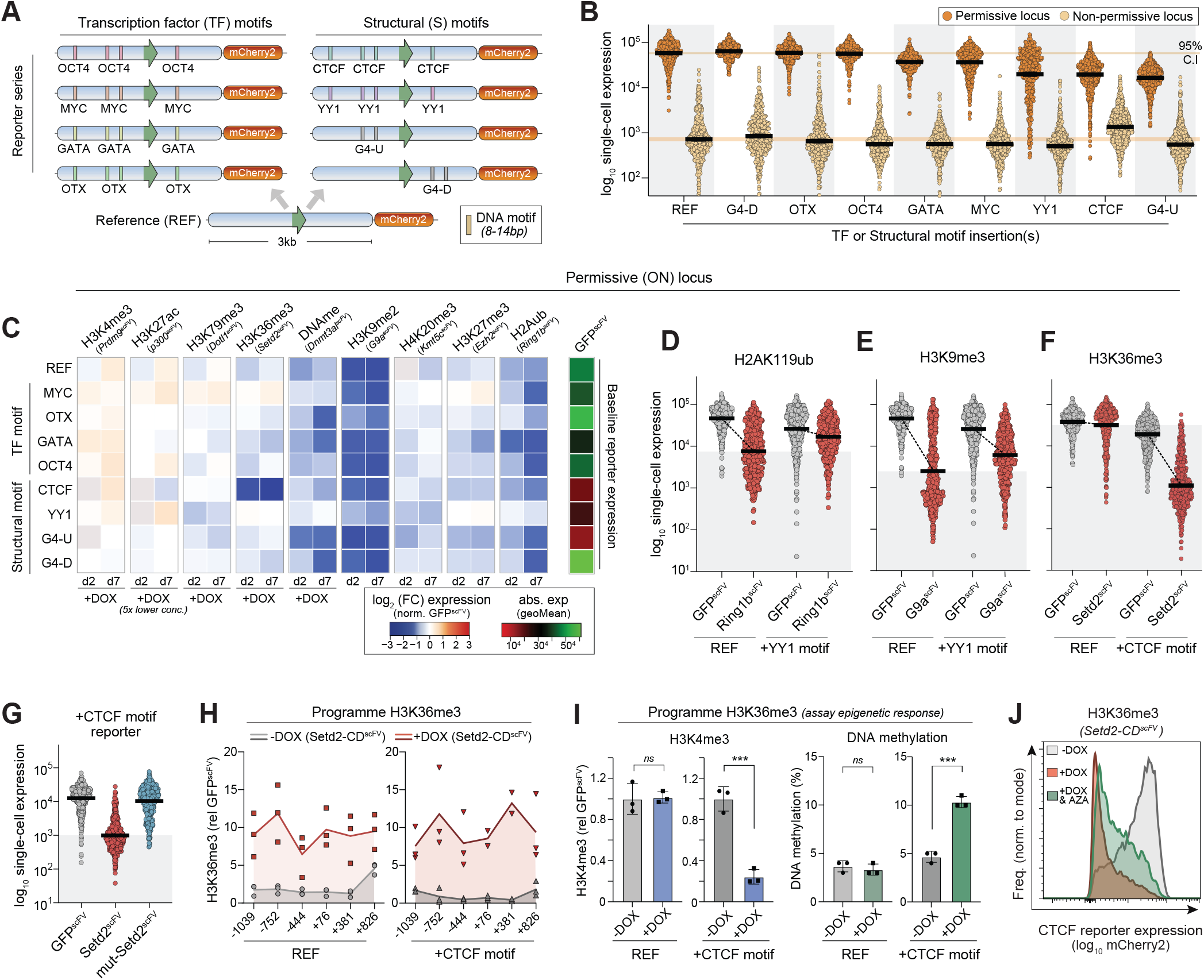
Functional interplay between chromatin marks and transcription factor motifs. **(A)** Schematic of the reporter series whereby each is identical apart from insertion of specific short sequence motifs. **(B)** Dot plots of mCherry2 expression from the indicated reporter type, integrated in either the permissive or the non-permissive locus. Each data point is a single cell and bars denote geometric mean. **(C)** Heat map showing the log^2^ fold-change in transcription at the ON locus upon programming the indicated chromatin mark (x-axis) to the indicated *cis* motif reporter (y-axis), relative to control GFP^scFV^ targeting. Data is shown after two days (d2) and 7days (d7) of DOX-induced epigenetic editing and corresponds to the average of four technical replicates. (**D-F)** Dot plots showing independent validations of functional interactions between programmed epigenetic marks and underlying sequence motifs. Each data point is log^10^ expression in a single cell carrying the indicated reporter, and bars denote geometric mean. (**G)** Dot plots of log_10_ single-cell expression of the +CTCF reporter after GFP^scFV^, Setd2^scFV^ (H3K36me3) or mut-Setd2^scFV^ targeting for 7 days. Bars denote the geometric mean. **(H)** Relative abundance of H3K36me3 at the reference (left) or +CTCF (right) reporters assayed by ChIP-qPCR before (-DOX) or after (+DOX) Setd2^scFV^ induction, across a ∼2kb region. Lines denote the mean of three replicates. (**I)** Bar plots showing the enrichment of H3K4me3 (left) and percentage of DNA methylation (right) on either the Reference or +CTCF reporters following programing of H3K36me3. Error bars represent S.D. with significance by unpaired t-test. **(J)** Representative flow cytometry plot showing expression of the +CTCF reporter before (-DOX) or after (+DOX) programming H3K36me3 +/the DNA methylation inhibitor AZA.

To systematically explore *cis* genetic x epigenetic functional interplays, we installed each chromatin modification, to each reporter, within each genomic context. We first focussed on the ‘ON’ reporter(s) (permissive locus), which as expected, were not significantly impacted by further deposition of positive marks H3K27ac, H3K4me3 and H3K79me2. In general, repressive modifications exhibited a good concordance in function across the reporter series. For example, H3K9me2/3 and H2AK119ub manifested strong silencing by day 7 (d7) of induction irrespective of most underlying motifs, with H3K9me2/3 exhibiting the faster repression kinetics (Fig 4C). Nevertheless, we did observe a number of striking functional interactions between specific marks and *cis* genetics, which were highly reproducible across replicates (Fig 4C & S8B-C). For example, the presence of YY1 motifs within an otherwise identical sequence effectively blocked the capacity for H2AK119ub and H3K27me3 to instruct transcriptional repression. Such YY1 sites also dampened the quantitative impact of DNA methylation and H3K9me2/3 (Fig 4C). Conversely, the presence of OTX motifs rendered the reporter more amenable to repression by DNA methylation. The most salient observation however related to H3K36me3, which generally has a weak and partially-penetrant effect on transcription across the series. However, programming H3K36me3 specifically on the +CTCF motif reporter resulted in a switch-like behaviour, with imposition of highly significant transcriptional silencing beyond levels obtained with any other repressive mark across any context (Fig 4C).

To validate these contextual relationships, we generated independent knock-in reporter lines and targeted them with specific chromatin modifications. We confirmed inclusion of *cis* YY1 motifs robustly neutralised the repressive activity of H2AK119ub and H3K9me2/3 (Fig 4D-E). Quantitatively this meant expression was diminished by only 1.5-fold and 4.3-fold by H2AK119ub and H3K9me2/3 respectively, rather than 6.1-fold and 18.5-fold repression of the baseline reporter that lacked 12bp YY1 sites. Whilst the link between DNA methylation and OTX motifs was variable (Fig S8C), we reproducibly observed that the inclusion of CTCF motifs, within an otherwise identical genomic context, licensed H3K36me3 to instruct transcriptional silencing exceeding 20-fold at the population level, with >98% of cells responding (Fig 4F & S8B). In contrast there is almost no effect of H3K36me3 on the REF promoter.

We confirmed that this context-dependent H3K36me3 activity is driven by the mark itself, since targeting the mut-Setd2-CD^scFV^ effector had no impact on transcription (Fig 4G). Moreover, H3K36me3 is programmed to comparable (high) levels on both the REF and the +CTCF motif reporter types, ruling out that differential responses are linked with disparities in initial epigenetic editing (Fig 4H). However, upon H3K36me3 programming specifically at the +CTCF reporter, the level of H3K4me3 decreased sharply. In contrast, H3K4me3 remained unaffected by *de novo* H3K36me3 on the REF (Fig 4I). DNA methylation was also preferentially increased only on the +CTCF reporter following installation of H3K36me3 (Fig 4I). Thus, equivalent levels of H3K36me3 induce different epigenetic cascades depending on the underlying genetic sequence/motifs.

To test the functional significance of this, we targeted Setd2-CD^scFV^ to the +CTCF reporter coincident with 5-azacytidine (AZA) treatment, a potent DNA methylation inhibitor. AZA reduced the fraction of cells that fully switch OFF the +CTCF reporter in response to H3K36me3, implying a partial role for DNA methylation recruitment downstream of H3K36me3 function (Fig 4J). We conclude the functional output of H3K36me3 is sensitive to the *cis* genomic sequence and its susceptibility to downstream epigenomic remodelling. Taken together, these data exploit a controlled system to reveal that underlying genetic motifs or variants mediate complex regulatory interactions with epigenetic modifications that quantitatively influence the transcriptional response. This implies the precise function of a chromatin modification ‘peak’ is not unequivocal, but highly context-dependent.

### Functional interplay between activating marks and TF motifs

We next examined genetic x epigenetic interactions within the transcriptionally silent genomic context, noting that with the exception of H3K9me2/3, and H3K36me3 on the CTCF reporter, repressive modifications could not drive further silencing irrespective of genetic motifs. Programming H3K79me2 installed weak activation, with no major variation across *cis* contexts. However, we observed significant interplay between H3K4me3- and H3K27ac-mediated activation and underlying sequence motifs. For example, the presence of either MYC or YY1 sequence motifs strongly reduced or even neutralized the activity of both H3K4me3 and H3K27ac (Fig 5A). Conversely, OCT4 and OTX2 motifs synergised with H3K4me3 and H3K27ac, respectively, potentiating their positive effect on transcription in ESC. We again validated our results by introducing the epigenetic editing machinery into independent knock-in reporter clones. This confirmed that the function of H3K27ac is reciprocally modulated by the presence of short motifs - MYC (attenuates) and OTX (enhances) (Fig 5B-C) - which manifests as a 1.4-fold versus 5.1-fold activation by H3K27ac, relative to 3.5-fold in the REF context. We further confirmed significant interactions between +MYC, +OCT4, or +CTCF *cis* contexts and H3K4me3 effects (Fig 5D & S8C).

**Figure 5.**
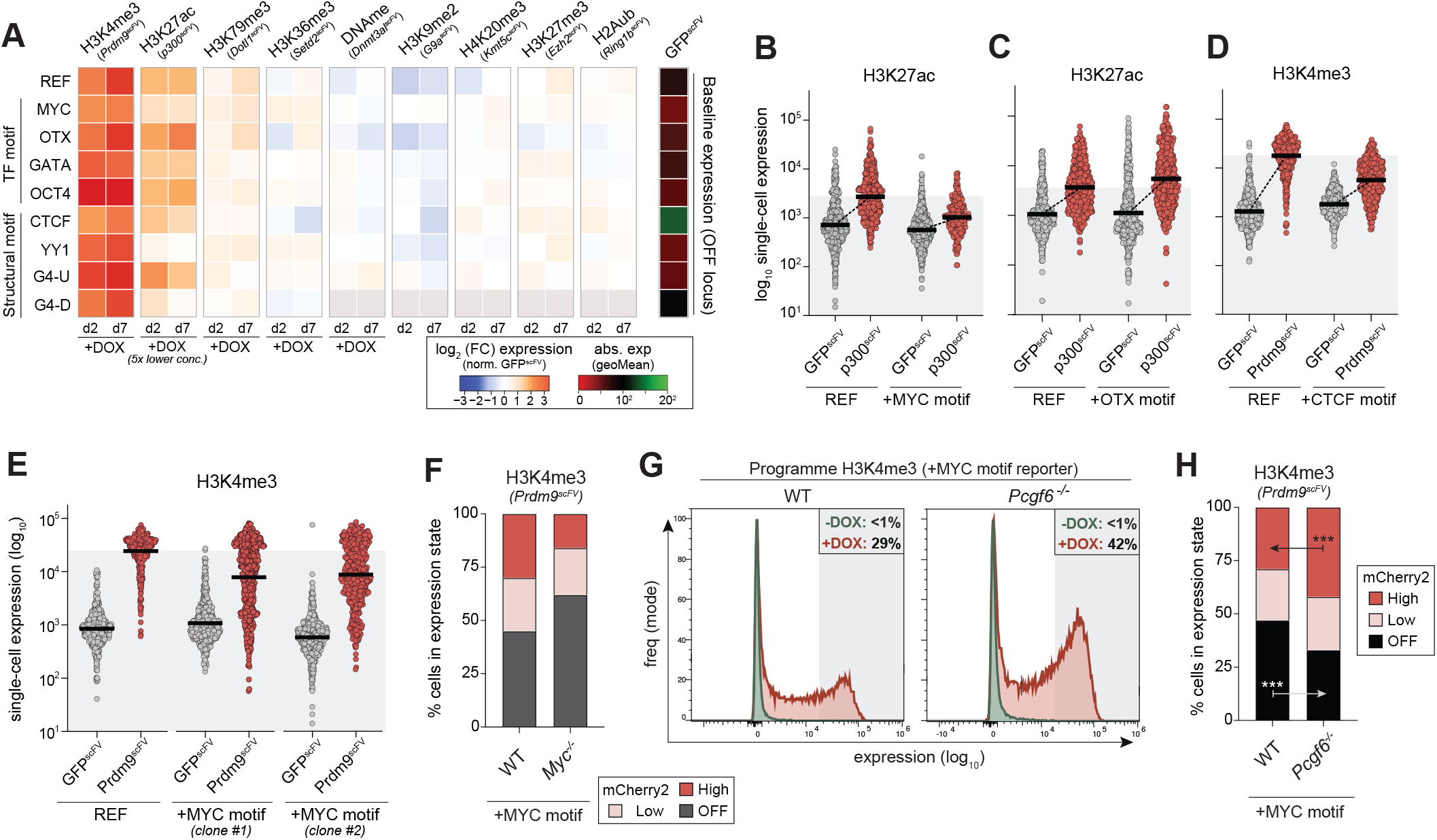
Instructive activity of chromatin modifications is throttled by *cis* genetics. **A)** Heat map showing the log_2_ fold-change in transcription at the OFF locus upon programming the indicated chromatin mark (x-axis) to the indicated *cis* motif reporter (y-axis), relative to control GFP^scFV^ targeting. Data is shown after two (d2) and seven days (d7) of DOX-induced epigenetic editing and corresponds to the average of four technical replicates. **(B-D)** Dot plots showing independent validations of functional interactions between programmed epigenetic marks and underlying sequence motifs. Each data point is log_10_ expression of the indicated reporter variant in a single cell after control GFP^scFV^ or specific CD^scFV^ epigenetic editing for 7 days. Bars denote the geometric mean. **(E)** Dot plots showing single-cell expression of independent +MYC reporters is limited after induction of H3K4me3, relative to control REF reporter. **(F)** Contingency plot indicating the fraction of cells that acquire a “off”, “low” or “high” expression state following H3K4me3 programming, in a wild-type (WT) or a *Myc*^-/-^ genetic background. **(G)** Representative flow cytometry plot showing the inactive +MYC reporter expression before (-DOX) or after (+DOX) Prdm9^scFV^ targeting for 5 days, in either a wild-type (WT) or a *Pcgf6*^-/-^ genetic background. **(H)** Contingency plot indicating an elevated fraction of cells acquire the “high” expression state from +MYC reporter following H3K4me3 programming in *Pcgf6*^-/-^ ESC. Significance is calculated by two-way ANOVA *\*P<0*.*05 **P<0*.*01, ***P<0*.*001*.

To investigate the mechanistic nature of such context-dependent responses, we focused on the attenuation of H3K4me3 (and H3K27ac) function by MYC motifs, which we observed across clones (Fig 5E). We first knocked out *Myc* in ESC carrying the +MYC reporter using CRISPR. Programming H3K4me3 in *Myc*^*-/-*^ ESC still led to attenuated activation of the +MYC motif reporter relative to REF, and indeed actually increased the fraction of non-responding (OFF) cells, potentially because *Myc* is associated with transcription activation ^45^. This implies that recruitment of *trans* MYC protein does not underpin the interaction between the *cis* DNA motif and H3K4me3 (Fig 5F). We therefore next focused on the variant polycomb complex PRC1.6, which also specifically binds MYC motifs (also known as E-box) ^46^. We generated knockout ESC lines for the key PRC1.6 component *Pcgf6*, and installed H3K4me3 at the +MYC reporter in *Pcgf6*^*-/-*^ cells. This reproducibly led to a rescue of attenuation, and significantly increased activation relative to programming H3K4me3 in WT controls (Fig 5G).

Specifically, whilst the fraction of cells weakly activating the +MYC reporter in response to H3K4me3 was similar between WT and *Pcgf6*^*-/-*^ cells, the fraction of cells that fully activated the reporter was significantly increased in the absence of *Pcgf6* (*P<0*.*001*; unpaired t-test) (Fig 5H). These data suggest that recruitment of PRC1.6 to promoters via MYC/E-box motifs provides a genetically encoded mechanism that limits the maximal expression induced by epigenetic systems such as H3K4me3. More generally, these data underline the relevance of genomic context in mediating the quantitative regulatory output of a chromatin mark.

### Naïve ESC antagonise epigenetic memory

We next deployed our editing toolkit to interrogate other regulatory questions. We first asked whether epigenetically programmed transcriptional states can be inherited through mitotic divisions and whether DNA context impacts this. We targeted each CD^scFV^ to each reporter in each genomic context for seven days to install the panel of epigenetic modifications, and then withdrew DOX to remove the inducing signal. Despite robust initial transcriptional responses, upon seven days withdrawal of the editing machinery (DOX wo) we observed no significant long-term memory of either activated or repressed reporter activity (Fig 6A-B). This was evident for all tested genetic contexts and regardless of genomic location, implying that transcriptional changes instigated by *de novo* chromatin marks are robustly reset to baseline in naïve ESC. Such lack of ‘epigenetic memory’ is consistent with recent observations that acquired heterochromatin domains do not propagate in naïve pluripotent cells ^38^.

**Figure 6.**
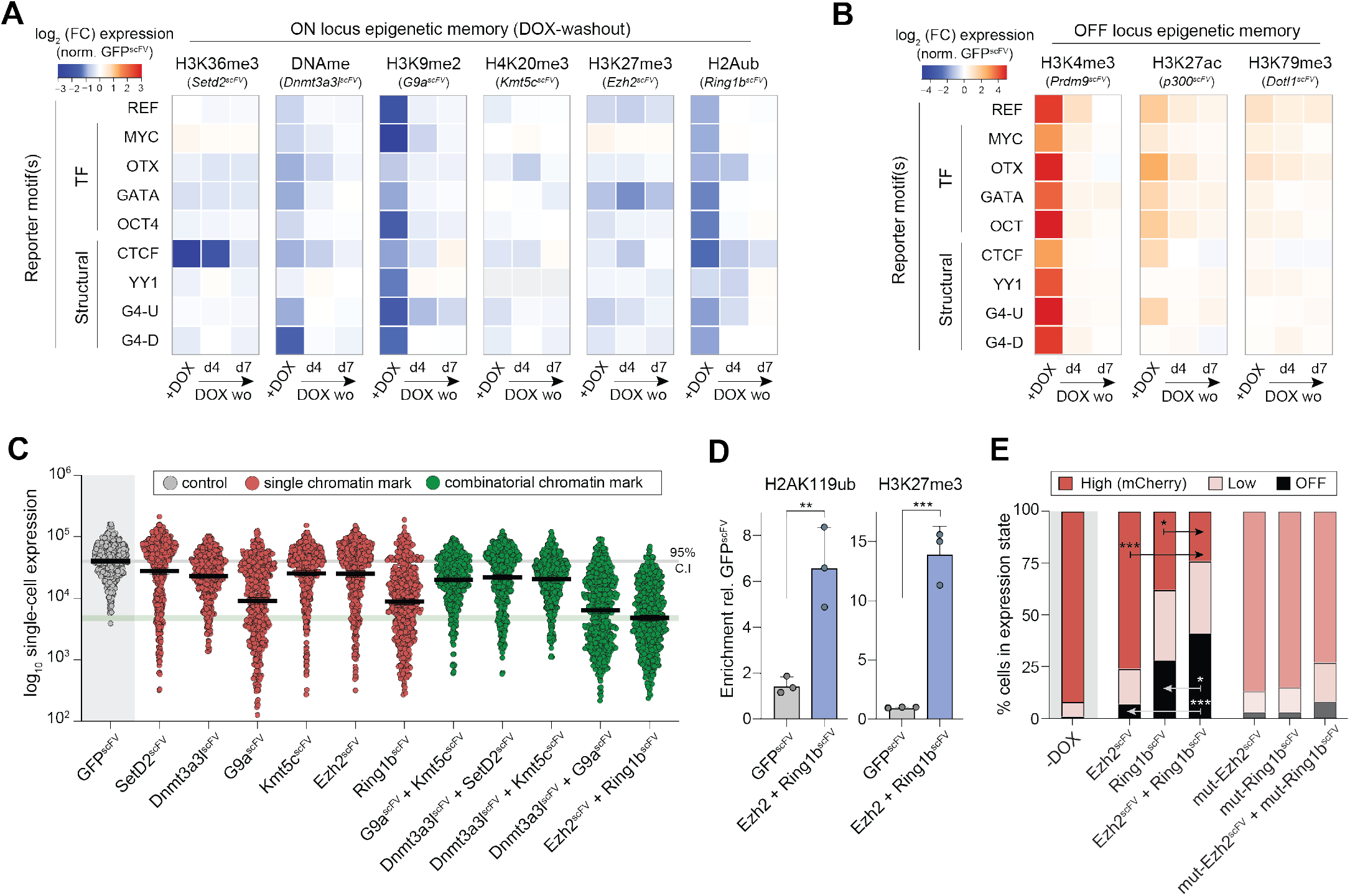
Combinatorial epigenetic editing reveal a functional synergy between H3K27me3 and H2Aub. **(A-B)** Heat map showing the log_2_ fold-change in transcription upon programming the indicated chromatin mark (x-axis) to the indicated motif reporter (y-axis), and then upon washout (DOX wo) for seven days to assay epigenetic memory. Shown areptranscriptional persistance effects at **(A)** the ON locus and **(B)** the OFF locus. **(C)** Representative dot plots indicating log_10_ expression after control GFPscFv, single CD^scFv^ or multiplex CD^scFv^ targeting for seven days to programme combinatorial marks. Each data point represents a single cell and bars denote geometric mean. **(D)** Bar plots showing enrichment of H2AK119ub (left) and H3K27me3 (right) on the ON Reference reporter assayed by CUT&RUN-qPCR following control GFP^scFv^ or combinatorial Ezh2^scFv +^ Ring1b scFv targeting. Shown is the mean of three biological replicates, error bars represent S.D, significance by unpaired I-test. **E**. Contingency plot indicating an elevated fraction of cells acquire the “OFF” expression state following combinatorial H3K27me3/H2AK119ub programming. Significance is calculated by two-way ANOVA *\*P<0*.*05 **P<0*.*01, ***P<0*.*001*.

### Combinatorial epigenetic editing reveals functional synergy of PRC1 and PRC2 activity

We finally asked if and to what extent combinatorial chromatin marks interact with one another to synergise or antagonise their quantitative effects on transcription. Our modular dCas9^GCN4^ system can recruit multiple CD^scFV^ effectors simultaneously. We therefore induced pairs of CD^scFV^ together, focusing on combinatorial marks that co-occur on chromatin (Fig 6C). Amongst functional interactions, we noted that concomitant deposition of H3K9me2/3 and DNA methylation (Dnmt3a3l-CD^scFV^ + G9a-CD^scFV^) increased the robustness of the transcriptional silencing response, relative to deposition of each mark singularly. Specifically, whilst the maximal level of repression amongst single cells was similar to H3K9me2/3, there was an increase in the fraction of cells that fully silenced expression when DNA methylation was co-targeted (35%±6 vs 41%±4), indicating these marks may cooperate to confer robustness (Fig 6C & S9A). Accordingly, when DNA methylation was inhibited following H3K9me2/3 deposition using AZA (Fig S9B), an elevated percentage of cells did not fully silence reporter activity (Fig S9C).

The most striking synergy however came from co-targeting H3K27me3 and H2AK119ub (Ezh2-CD^scFV^+ Ring1b-CD^scFV^), which instigated a highly significant increase in the single-cell penetrance of silencing, relative to installing either mark individually (Fig 6C-E & S9D-E). This effect was particularly intriguing since it is not clear whether the transcriptional impact of PRC1 (H2AK119ub) and PRC2 (H3K27me3) at polycomb domains arises from the sum of their individual activities. We confirmed that significant levels of both H3K27me3 and H2AK119ub are programmed by combinatorial targeting (Fig 6D). Moreover, independent ESC lines supported that such multiplex epigenetic editing led to a functional synergism, with 41% (±7% S.D) of cells reaching the fully OFF state, relative to deposition of H2AK119ub (28% ±7; *P=0*.*029*) or H3K27me3 (7% ±3; *P<0*.*001*) alone (Fig 6E & S9E. Importantly, catalytic mutant effectors registered only a subtle negative effect on reporter activity. Overall, these data suggest that combinatorial chromatin modifications can increase the single-cell penetrance of transcriptional responses, with H3K27me3 and H2AK119ub together exemplifying effects greater than the sum of their parts. Such functional interactions between marks provides an additional layer of context-dependency, and further uncovers the parameters that modulate the quantitative effects of chromatin modifications.

## DISCUSSION

The extent to which specific chromatin modifications are causative or consequential of DNA-templated processes, and in which contexts, is an area of intense debate^35,40^. To address this, we developed a comprehensive epigenetic editing toolkit that enables *de novo* installation of a repertoire of nine key chromatin marks at precise genomic loci with high efficiency. We leverage this platform to capture that acquisition of each tested modification is sufficient to trigger at least some transcriptional response, in at least some contexts, with overall effects ranging from non-functional through to order-of-magnitude expression changes. The precise quantitative impact and single-cell penetrance of a mark is contingent on multiple contextual factors however, and we provide direct evidence that underlying TF motifs, genomic positioning and combinatorial modifications interact to modulate the overall expression output. Thus, whilst our data imply that chromatin marks have the potential to causally instruct transcription programmes, they also highlight they represent one regulatory layer within multiple nonlinear governing mechanisms.

Amongst our findings we charted a function for H3K4me3, which is an evolutionary conserved marker of transcriptionally active promoters, and directly recruits the preinitiation complex (PIC) via TAF3 ^47,48^. Nevertheless, loss-of-function studies across model systems suggest that H3K4me3 is not required for the majority of gene expression^14,49,50^. Indeed, recent studies have implied that promoter H3K4me3 primarily reflects a consequence of transcription activity^15^. However, we report that *de novo* acquisition of H3K4me3 can instruct robust transcriptional upregulation from a subset of silent promoters. We confirm the direct effect of the mark *per se* using an array of different H3K4me3 programming tools, catalytic-mutant controls, and *Mll2*^*CM/CM*^ ESC that specifically lack H3K4me3. The cumulative studies point toward a dual-feedback relationship whereby transcription itself promotes downstream accumulation of H3K4me3, but reciprocally, that *de novo* acquisition of H3K4me3 can trigger transcription. Indeed, H3K4me3 appears important for the timely activation of gene subsets during pluripotent state transition here, and in germline specification^51^.

Mechanistically, we find that programming H3K4me3 initiates an epigenetic cascade that includes extensive promoter acetylation, which is required for the functional impact of H3K4me3. This is likely reinforced, to some extent, by the transcription machinery having direct affinity for H3K4me3^47,52^. Nevertheless, it is important to note that H3K4me3 activity is contingent on appropriate TF in the cellular milieu and indeed, only a fraction (∼35%) of silent genes responded to *de novo* H3K4me3. In this respect, acquisition of H3K4me3 may instruct transcriptional upregulation primarily by antagonising epigenetic repression, thereby establishing a permissive environment for relevant TF. Such a model is consistent with loss-of-function studies showing H3K4me3 depletion can be rescued by concomitant depletion of H3K27me3 and DNA methylation. Indeed, programming H3K4me3 here directly evicted H3K27me3, whilst concurrently driving H3K27ac enrichment. Notably, whilst we observed major gene upregulation (>50-fold) following *de novo* H3K4me3, previous studies have reported programming H3K4me3 either has subtle effects (typically <2-fold) ^26^, or no measurable impact ^34^. This difference may be rooted in the efficiency of H3K4me3 editing, with our optimised toolkit amplifying the magnitude, and particularly the genomic breadth, of *de novo* H3K4me3 domains. Indeed, gene expression levels are tightly correlated with both the intensity and breadth of promoter H3K4me3 ^53^, and we observed dose-dependent transcriptional responses to epigenetic editing. Taken together, we propose that sufficient *de novo* H3K4me3 can antagonise extant repressive mechanisms and enable transcription initiation, if appropriate trans-acting factors are present.

Transcription factors sit at the apex of transcriptional regulation cascades, and therefore focus on the role of chromatin modifications has often fallen on how they directly or indirectly modulate TF activity. This is evident for DNA methylation for example, which impairs TF such as NRF1 and BANP from binding cognate sites ^54,55^, and histone modifications, which can impede non-pioneer TF activity ^36,56^. Less is understood about the reciprocal relationship, whereby TFs modulate the functional output of a chromatin modification. By quantifying the instructive potential of multiple marks, we were subsequently able to use reductionist strategy to dissect how underlying DNA sequence or TF motifs influence such chromatin function to tune outputs. For example, the presence of YY1 motifs limited the repressive potential of both H3K9me2/3 and polycomb marks, effectively conferring partial resistance to epigenetic silencing. Reciprocally, MYC/E-box motifs restricted activation by *de novo* H3K4me3 or H3K27ac. This reflects the activity of the PRC1.6 complex that occupies E-box motifs ^46^, which in turn therefore act as genetically-encoded signals that threshold maximal activation. Such *cis* genetic x epigenetic interplays that shape the expression space could have implications for the evolutionary potential of gene regulatory networks ^57^.

The most striking interaction entailed a switch-like behaviour of H3K36me3, which instructed strong reporter silencing only in the context of *cis* CTCF motifs. Such context-dependent H3K36me3 function could be linked with CTCF-mediated nucleosome phasing, 3D looping, direct transcription modulation and/or chromatin insulation ^43^, which necessitates future study. More generally, understanding the bidirectional regulatory relationship(s) between the genome and epigenome is key towards deciphering how DNA sequence variants influence phenotypic traits ^58^. For example, a given sequence variant that alters TF binding, thereby creating an expression quantitative trait locus (eQTL), may be unmasked or neutralized depending on the interactions with the overlying epigenetic modification(s). A further contextual parameter relates to the interplay between overlapping chromatin modifications. We find combinatorial H3K27me3 and H2AK119ub marks synergise to enhance the fraction of responsive cells, but not absolute repression. Such epigenetic ‘penetrance’ effects at the single-cell level also contributed to differential responses to singleplex epigenetic editing. This implies there is a equilibrium of regulatory forces at steady state, with programming of more influential (or combinatorial) marks having a greater, but not unequivocal, probability of overcoming the governing *status quo* in each cell. Importantly however, whilst our data imply that chromatin marks can be instructive, they also emphasize that impacts are context-dependent. This argues against a hard-wired ‘histone code’ whereby specific patterns of chromatin marks elicit a specific output, and instead points toward a nonlinear regulatory network that produces quantitative outputs depending on myriad inputs including TF binding, chromatin architecture, *cis* genetics, metabolic state, and indeed epigenetic modifications themselves.

In summary, our study captures the principles of how *de novo* chromatin modifications can causally influence gene expression across contexts. Moreover, the modular epigenetic editing toolkit provides a framework to explore regulatory mechanisms across DNA-templated processes, and to precisely manipulate chromatin for desirable responses in disease models.

## METHODS & MATERIALS

### Cell culture

Wildtype mouse embryonic stem cells (mESCs) were derived freshly (mixed 129/B6, XY) and cultured on gelatin-coated cell culture plates under naïve conditions (2i/LIF). Routine passaging was performed in N2B27 basal culture medium (NDIFF, Takara #y40002), supplemented with 1 μM PD0325901 and 3 μM CHIR99021 (both from Axon Medchem), 1,000 U/ml leukaemia inhibitory factor (LIF; in house production), 1% FBS (Millipore) and 1% penicillin/streptomycin (Gibco). All culture media was filtered through a 0.22μm pore Stericup vacuum filtration system (Millipore). Cells were maintained at 37°C in a 5% CO_2_ humidified atmosphere and were passaged every 2 days by dissociation with TrypLE (Thermo Fisher Scientific). Culture media was replaced with fresh stocks daily. Mycoplasma contamination was tested routinely by ultrasensitive qPCR assay (Eurofins).

### Generation of reporter cell lines

We designed a Reference reporter to provide a baseline context, and to enable the influence of subsequently inserting sequence motifs or variants to be assessed. We used the endogenous EF1α core promoter (∼200bp) embedded into a DNA sequence context selected from human chromosome 7 (chr7:41344065-41346105, GRCh38/hg38) to be neutral in respect of genomic features, including: depleted of transcription factor motifs, GC percentage (50%), lacking retrotransposons, and without epigenetic enrichments. The resulting cassette (∼3kb) was designed as a gBlock gene fragment from Integrated DNA Technologies (IDT), and amplified by PCR using Q5 hot start high-fidelity polymerase (NEB #M0494S) and primers with appropriate overhangs. This was inserted by In-fusion HD-Cloning into a recipient vector upstream of a Kozak sequence, the mCherry2-H2B fluorescence coding sequence, and a poly-A motif. The assembled reporter construct (DNA::EF1α Pr::DNA::mCherry2-H2B::pA) was sequence-verified, and then PCR amplified with Q5 polymerase, using ultramer DNA oligos (Eurofins) carrying 200bp-long overhangs homologous to DNA sequences flanking the desired genomic insertion site(s). Specifically, we chose two intergenic genomic insertion sites that differentially support transcription. Firstly, a permissive landing site (chr9:21545329; ON locus, TIGRE) and secondly a non-permissive landing site that only supports weak transcription (chr13:45253722; OFF locus), albeit within a euchromatic domain.

To insert the cassettes into each locus, we transfected 1μg of PCR-amplified dsDNA reporter into naïve mESCs together with spCas9 plasmid pX459 (*Addgene #62988*), carrying a single gRNA complementary to the genomic integration site. After puromycin selection (1.2ug/ml) for transient px459 transfection (2 days), mCherry2 positive cells that were candidates for correct insertion were purified by fluorescence-activated cell sorting (FACS). Single clones were expanded and correct monoallelic (hemizygous) integration of the reporter was verified by PCR genotyping and Sanger Sequencing (Azenta). The full allelic series of reporter variants, which each comprised the same baseline sequence as the Reference, but with insertion of several discrete transcription factor or structural motifs (see Supplementary materials for more info) were also ordered as gBlock Gene Fragments from IDT. Generation of the complete reporter cassette and genomic integration was carried out as described above for the Reference to generate a total of eighteen independent reporter lines (nine reporter variants in two genomic locations), each with independent clones. We validated independent insertions of each reporter to confirm reproducibility.

### Generation of epigenetic editing toolkit constructs

Epigenetic editing tools comprising a nuclease dead (d) Cas9^GCN4^ and the catalytic core of chromatin-modifying enzymes were cloned into PiggyBac recipient plasmids by homology arm recombination using In-fusion HD cloning (Takara #639650). Specifically, the *Streptococcus Pyogenes* dCas9^GCN4^ was PCR amplified from the PlatTET-gRNA2 plasmid ^37^ (Addgene #82559), and sub-cloned under the control of a DOX-inducible TRE-3G promoter into a PiggyBac backbone. The vector also carries the TET-ON 3G transactivator and hygromycin resistance.

For all chromatin-modifying ‘effector’ plasmids, the scFV domain and a superfolder (sf)GFP coding sequence were amplified from the PlatTET-gRNA2 plasmid (Addgene #82559) and fused in frame with the catalytic domain (CD) or the full-length (FL) of mouse *Prdm9, P300, Dot1L, G9a, Kmt5c, Setd2, Ezh2 and Ring1b*, all amplified from early passage ESC cDNA. *Dnmt3a* CD and the C-terminal part of mouse Dnmt3L (3a3L) were amplified from pET28-Dnmt3a3L-sc27 (Addgene #71827). The resulting constructs (collectively: CD^scFV^) were cloned in PiggyBac recipient vectors under the control of the TRE-3G promoter. These vectors also carry constitutive expression of a Neomycin resistance gene. The control GFP^scFv^ effector was cloned as described above but lacks any chromatin modifying domain. Finally, catalytic mutant (mut-CD^scFV^) effectors were also cloned as described above. Specific mutations that abolish the catalytic activity of each CD^scFV^ but that retain protein stability were introduced during PCR amplification with oligonucleotide primers designed with precisely mismatched nucleotides. The catalytically-inactivating point mutations introduced in each CD^scFV^ are: Prdm9, G282A; p300, D1398Y; Dot1l, GS163-164RC; Setd2, R1599C; Dnmt3a C706S; G9a, Y1207del; Kmt5c, NHDC182-185AAAG; Ezh2, Y726D; Ring1b, I53S; Set1a, S1631I.

The guide RNA plasmid, carrying an enhanced gRNA scaffold ^39^, was amplified from Addgene plasmid #60955 and cloned into a PiggyBac recipient vector, which also constitutively expressed puromycin resistance and TagBFP. All gRNAs used to target the epigenetic editing system were designed using the GPP web portal (Broad Institute). gRNA forward and reverse strands carrying appropriate overhangs (10 μM final concentration) were annealed in buffer containing 10 mM Tris, pH 7.5–8.0, 60 mM NaCl, 1 mM EDTA, at 95°C for 3 min and allowed to cool down at RT for > 30 min. Annealed gRNAs were ligated with T4-DNA ligase (NEB #M0202S) for 1 h at 37°C into the PiggyBac recipient vector previously digested with BlpI (NEB #R0585S) and BstXI (NEB #R0113S) restriction enzymes. Final plasmids were amplified by bacteria transformation and purified by endotoxin-free midi-preparations (ZymoResearch #D4200). Correct assembly and sequences were confirmed by Sanger sequencing (Azenta).

### Epigenetic editing assays

For stable integration of the epigenetic editing system, mESC lines were co-transfected with dCas9^GCN4^, one or more CD^scFV^ (or control GFP^scFV^), and gRNA plasmids in addition to the PiggyBac transposase vector using 10:20:2:1 molar ratio, respectively. Cells with successful integration of all three constructs were enriched by successive antibiotic selection with hygromycin (250 μg/ml) for 5 days, neomycin (300 μg/ml) for 5 days and puromycin (1.2 μg/ml) for 2 days. After allowing cells to recover and expand, expression of dCas9^GCN4^ and CD^scFV^ was induced by supplementing the culture media with doxycycline (DOX) (100 ng/ml) for either 2 or 7 days, with the exception of p300-CD^scFV^, whereby we used 5ng/ml DOX to mitigate against OFF-targeting. Correct induction of all epigenetic editing components results in double GFP and BFP positive cells (GFP+; BFP+). Activity of endogenous target genes or reporter (mCherry2) was analysed by quantitative PCR or quantitative flow cytometry, by sorting/gating for analysis only GFP^+^; BFP^+^ cells, which have correctly induced the editing system (typically >75% cells). For experiments employing the p300 inhibitor A485, cells were stimulated with 100ng/ml DOX for 3 days and, in parallel treated with 3μM A485 (Cayman Chemical, #24119). Where indicated 1μM 5-azacytidine (AZA, from Sigma-Aldrich) was included in media and replaced daily for 3 days in a row.

For epigenetic memory experiments, cells were washed thoroughly with PBS, and subsequently cultured in the absence of DOX, which led to a rapid downregulation of the epigenetic editing machinery (GFP-). Memory of reporter expression changes was quantified by flow cytometry after 4 or 7 days of DOX washout (DOXwo) in cells that were confirmed to have fully switched off the epigenetic editing tool (BFP^+^/GFP^-^ cells; typically >99%).

### Transfection

DNA transfection was performed with Lipofectamine 3000 (Thermo Fisher Scientific #L30000015). Cells were seeded one day in advance so as to reach ∼60% confluency on the day of transfection. Appropriate amounts of DNA were calculated according to manufacturer’s instructions. Media were changed after 8 h, and replaced with fresh antibiotic containing medium.

### Generation of genetically edited ESC lines

Knockouts (KO) cell lines for *Pcgf6* and *Myc* were generated by means of CRISPR/Cas9 genome editing. Specifically for each target gene, two plasmids (pX459) were transiently transfected into low-passage wild-type ESCs that had previously been engineered to carry a specific knock-in reporter. Each plasmid expressed one of two gRNAs targeting the flanking introns of a critical coding exon in the gene of interest (*c-Myc, Pcgf6*) (see table of gRNAs) and Cas9. Critical exons were present within all known isoforms and gRNAs were designed with the goal of specifically deleting the entire exon. After transfection, cells were selected with puromycin (1.2 μg/ml) for 3 days and subsequently seeded at low density (1,000 cells per 10cm^2^) for single clone isolation. Following expansion, single clones were screened for homozygous genetic editing by PCR genotyping (see table of primers) and dual loss-of-function (frameshifted) alleles were confirmed by Sanger sequencing (Genewiz). For generation of precision edited catalytic-mutant *Mll2 (Mll2*^*CM/CM*^) lines, homozygous ESC were freshly derived from heterozygous FVB crosses carrying an *Mll2* Y2602A mutant allele.

### Flow cytometry

Cells were washed with PBS and gently dissociated into single-cell suspension using TrypLE, followed by resuspension in FACS buffer comprised of PBS with 1% FBS, and filtered through a 40μm cell strainer (BD, cup-Filcons #340632). A FACS Aria III (Becton Dickinson) or Attune NxT Flow Cytometer (Thermo Fisher Scientific) were used for sorting or analysis, respectively. 96-well plates containing the different combinations of reporter x epigenetic effector cell lines were analyzed using the Attune NxT Flow Cytometer Autosampler and resulting data was used to generate the heat maps shown in Fig. 4C and 5A. Alternatively, specific reporter x epigenetic effector cell lines were generated and cultured in 12 well plates and samples were analyzed one by one using the single sample line of the Attune NxT Flow Cytometer. Flow cytometry data analysis was performed with FlowJo v10.5.3 (Tree Star, Inc.).

To generate dot plots shown in this study, the FlowJo software was used first to gate for live cells and then for cells expressing all epigenetic editing components (GFP^+^; BFP^+^). The resulting population was randomly down-sampled to 1000 cells. The mCherry2 scaled fluorescent values corresponding to the relative expression intensities for each cell were exported, and imported into Prism GraphPad statistical software. Dot plots were constructed with the geometric mean of the raw data shown (black bar). For dot plots representative of the individual reporter expression, prior to transfection of the editing machinery (Fig 4B), analysis was performed as described above, except that no GFP^+^; BFP^+^ gating was performed and mCherry2 single cell values were obtained from the whole population of live cells. To generate histograms, the parental GFP^+^; BFP^+^ cell population was selected as above and the frequency distribution of the flow data was plotted versus mCherry2 fluorescence intensity using a log_10_ scale. The bisector gating tool was then used to split histograms in two sectors corresponding to mCherry2 ON expression state and mCherry2 OFF expression state, based on negative and positive controls. Alternatively, the ranged gate tool was used to split the histogram in three sectors corresponding to mCherry2 “high”, mCherry2 “low” and mCherry2 “OFF” expression states. Identical gates were applied to all samples within an experiment.

Finally, to generate the heat maps, the mCherry2 scaled fluorescent values for 1000 GFP^+^; BFP^+^ cells were obtained and the geometric mean for each sample (indicating reporter expression after GFP^scFV^ or specific CD^scFV^ effector targeting) was calculated. The geometric mean of each CD^scFV^ effector was normalized against the corresponding geometric mean of GFP^scFV^ to obtain the fold change of reporter expression following epigenetic editing (geometric mean CD^scFV^ effector/geometric mean GFP^scFV^). The normalised geometric mean values coming from four technical replicates of the experiments were averaged and log_2_ transformed. Log_2_ fold-change values were plotted in R.

### RNA extraction, library preparation and sequencing

Total RNA was extracted from cells using the Monarch Total RNA Miniprep Kit (NEB #T2010), following manufacturer instructions. Purified RNA was quantitated with a Qubit Fluorometer (Thermo Fisher Scientific) and quality checked with an automated electrophoresis system (Agilent Tape Station system) to ensure RNA integrity (RIN >9). Precisely 1μg of each RNA sample was used to prepare sequencing libraries using the NEBNext Ultra II directional RNA library kit by the EMBL Genomics facility. Libraries were sequenced on the Nextseq Illumina sequencing system (paired-end 40 sequencing). Raw Fastq reads were trimmed to remove adaptors with TrimGalore (0.4.3.1, -phred33– quality 20–stringency 1 -e 0.1–length 20), quality checked and aligned to the mouse mm10 (GRCm38) genome using RNA Star (2.5.2b-0, default parameters except for–outFilterMultimapNmax 1000). Analysis of the mapped sequences was performed using Seqmonk software (Babraham bioinformatics, v1.47.0) to generate log_2_ reads per million (RPM) or gene length-adjusted (reads per kilobase million, RPKM) gene expression values. Differentially expressed genes (DEG) were determined using the DESeq2 package (v.1.24.0), inputting raw strand-specific mapping counts and applying a multiple-testing adjusted (FDR) *P< 0*.*05* significance threshold, and log_2_ fold-change filter where indicated.

### RT-qPCR

Total RNA was extracted from cells using the Monarch Total RNA Miniprep Kit (NEB #T2010), following manufacturer instructions. After quantification using a Qubit Fluorometer (Thermo Fisher Scientific), 1μg of each sample was DNAase treatment, and inputted into cDNA synthesis by incubation with a mixture of random hexamers and reverse transcriptase (TAKARA PrimeScript RT Reagent Kit with gDNA Eraser, Takara Bio #RR047A). The resulting cDNA was diluted 1:10 and 2 μl of each sample was amplified using a QuantStudio 5 (Applied Biosystems) thermal cycler, employing the SYgreen Blue Mix (PCRbio) and pre-validated gene-specific primers that span exon-exon junctions. Results were analyzed using 2–ΔΔCt (relative quantitation) with the QuantStudio 5 software and normalized to the housekeeping gene *Rplp0*.

### Bisulphite pyrosequencing

DNA bisulfite conversion was performed starting from a maximum of 1 × 10^5^ pelleted cells per sample using the EZ DNA Methylation-Direct kit (Zymo Research #D5021), and following the manufacturer’s instructions. Target genomic regions were amplified by PCR using 1μl of bisulfite-converted DNA and specific primer pairs, one of which is biotin-conjugated, using the PyroMark PCR kit (Qiagen #978703). 10μl of the PCR reaction was used for sequencing using the dispensation orders (below) generated by the PyroMark Q24 Advanced 3.0 software, along with PyroMark Q24 advanced reagents (Qiagen, #970902) according to manufacturer’s instructions. Briefly, the PCR reaction was mixed with streptavidin beads (GE Healthcare #17-5113-01) and binding buffer, denaturated with denaturation buffer using a PyroMark workstation (Qiagen) and released into a PyroMark Q24 plate (Qiagen) pre-loaded with 0.3μM of sequencing primer. Annealing of the sequencing primer to the single-strand PCR template was achieved by heating at 80°C for 2 min and cooling down at RT for 5 min. Pyrosequencing was run on PyroMark Q24 advanced pyrosequencer (Qiagen). Results were analysed with PyroMark Q24 Advanced 3.0 software.

#### Dispensation orders

##### Reference reporter

AGTGATCGTATACTAGTATAGAGATGTCGTGTAGTCTGTAGTGTAGATGTCGTATGATCGTATATGTTCTGA

##### Col16a1

ATCATCGATCTATCTCTACTAGTACATCGACATCGATATCGATCGACACACTCACATCGACTACTACAACTATCAGATCGACC

##### Hand1

CACTACGATAGCACTATCGACACATCATCACATCATCACACTCACATCGATCGACACCATACTCATCAGACTC

### CUT&RUN

The CUT&RUN (Cleavage Under Targets and Release Using Nuclease) protocol ^59^ was used to detect genomic enrichment of histone modifications. From 1×10^5^ to 1×10^6^ cells (depending on the selected antibody) were pelleted at 300g for 3 min following flow sorting. Cells were washed twice in Wash buffer (1 ml 1 M HEPES pH 7.5, 1.5 ml 5 M NaCl, 12.5 μL 2 M Spermidine, final volume brought to 50 ml with dH2O, complemented with one Roche Complete Protease Inhibitor EDTA-Free tablet). Pellets were then re-suspended in 1 ml of Wash Buffer and 10 μL of concanavalin beads (Bangs Laboratories #BP531-3ml) in 1.5ml Eppendorf tubes and allowed to rotate at RT for 10 min. Supernatant was removed by placing the samples on a magnet stand and 300μl of Antibody buffer (Wash buffer supplemented with 0.02% Digitonin and 2mM EDTA) containing 0.5-3 μg of target-specific antibody was added. Samples were left to rotate overnight at 4°C. Antibodies used were: Rabbit anti-H3K4me3 (Diagenode Cat#C15410003), Rabbit anti-H3K27me3 (Millipore Cat#07-449), Rabbit anti-H3K9me3 (Abcam Cat#ab8898), Rabbit anti-H2Aub (Lys119) (CST Cat#8240), Rabbit anti-H3K36me3 (Diagenode Cat#C15410192), Rabbit anti-H3K36me3 (Active Motif Cat#61101), Rabbit anti-H3K27ac (Active Motif Cat#39133), Rabbit anti-H3K79me2 (Abcam Cat#ab3594), Rabbit anti-H4K20me3 (Abcam, Cat#ab9053)

The following day, each tube was placed on a magnetic stand and cell-bead complexes were washed twice with cold Dig-wash buffer (Wash buffer containing 0.02% Digitonin), then re-suspended in 300μl of cold Dig-wash buffer supplemented with 700 ng/ml of purified protein-A::MNase fusion (pA-MNase). Samples were left to rotate on a rotor at 4°C for 1 h. After two washes in cold Dig-wash buffer cell-bead complexes were re-suspended gently in 50 μl of Dig-wash buffer and placed on an aluminium cooling rack on ice to be precooled to 0°C. To initiate pA-MNase digestion, 2 μl of 100 mM CaCl_2_ was added, samples were flicked to mix and immediately returned to the cooling rack. Digestion was allowed to proceed for 30 min and was then stopped by addition of 50 μl 2XSTOP buffer (340 mM NaCl, 20 mM EDTA, 4 mM EGTA, 0.02% digitonin, 250 μg of RNase A, 250 μg of glycogen). Samples were incubated at 37 °C for 10 min to release CUT&RUN fragments from the insoluble nuclear chromatin and centrifugated at 16,000g for 5 min at 4°C. The supernatant was isolated by means of a magnetic stand and transferred into a new tube while the cell-bead complexes were discarded. 2μl of 10% SDS and 2.5 μl of Proteinase K was added and the samples were incubated for 10 min at 70°C. Purification and size selection of DNA were performed using SPRI beads (Beckman Coulter #B23318) following the manufacturer’s instruction for double size selection with 0.5× and 1.3× bead volume-to-sample volume ratio. Purified DNA was eluted in 30 μl of Ultrapure water.

For analysis of specific genomic targets, CUT&RUN DNA fragments were subjected to quantitative qPCR analysis. A 1:10 dilution was performed and 2μl of diluted DNA was amplified by mean of a QuantStudio 5 (Applied Biosystems) thermal cycler using the SYgreen Blue Mix (PCRbio) and specific primers for both targeted and control genomic regions. Relative abundance of histone marks was determined by calculating the 2^-Ct value for each genomic region of interest and normalizing it against the 2^-Ct value of a positive control genomic locus (2^-Ct targeted region/2^-Ct positive control region). Data is then shown as relative fold change between experimental samples and control samples (e.g. CD^scFV^ over GFP^scFV^) with a randomly selected control replicate set as the baseline (=1).

For genome-wide analysis, CUT&RUN was performed as described above followed by library preparation. Specifically, eluted DNA fragments were purified and subject to size selection of DNA using SPRI beads (Beckman Coulter #B23318) following the manufacturer’s instruction for double size selection with 0.5× and 1.3× bead volume-to-sample volume ratio. Purified DNA was eluted in 30 μl of Ultrapure water and 10ng was inputted into the NEBNext Ultra II DNA Library Prep Kit for Illumina (NEB #E7645S) using the following PCR programme: 98°C 30 s, 98°C 10 s, 65°C 10 s and 65°C 5 min, steps 2 and 3 repeated for 12–14 cycles. After quantification and quality check with an automated electrophoresis system (Agilent Tape Station system), library samples were sequenced on the Nextseq Illumina sequencing system (paired-end 40 sequencing). Raw Fastq sequences were trimmed to remove adaptors with TrimGalore (v0.4.3.1, -phred33 --quality 20 --stringency 1 -e 0.1 --length 20), quality checked and aligned to the mouse mm10 genome with the inserted mCherry reporter using Bowtie2 (v2.3.4.2, -I 50 -X 800 --fr -N 0 -L 22 -i ‘S,1,1.15’ --n-ceil ‘L,0,0.15’ --dpad 15 --gbar 4 --end-to-end -- score-min ‘L,-0.6,-0.6’). Analysis of the mapped sequences was performed using seqmonk software (Babraham bioinformatics, v1.47.0) by enrichment quantification of the normalised reads. To identify promoters with H3K4me3 change in *Mll2*^*CM/CM*^, a 1kb window centered on the TSS was quantified amongst replicates and a normalised log fold-change (FC) filter applied between samples. Metaplots over genomic features were constructed by quantifying 100bp bins centered on the features of interest and normalised cumulative enrichments plotted.

### Chromatin immunoprecipitation-qPCR

3×10^6^ cells were dissociated with TryplE, resuspended in PBS and pelleted at 200g for 4 min at RT. After, PBS was removed and cell pellet was fixed in 1ml of 1% PFA for 10 min at RT, followed by centrifugation at 200g for 4 min. The supernatant was discarded and fixation was quenched by addition of 1ml 0.125 M glycine for 5 min at RT. Glycine was removed and pellets were washed twice with cold PBS. Samples were kept on ice from this stage onwards. Cells were resuspended in 1ml of cold Lysis buffer (50 mM HEPES pH 8.0; 140 mM NaCl; 1 mM EDTA; 10% glycerol; 0.5% NP40; 0.25% Triton × 100), incubated on ice for 5 min and subsequently spun down at 1200g for 5 min at 4°C. One wash in Rinse buffer (10 mM Tris pH 8.0; 1 mM EDTA; 0.5 mM EGTA; 200 mM NaCl) was performed, followed by another centrifugation at 1200g for 5 min at 4°C. Cell nuclei were then resuspended in 900 μl of Shearing buffer (0.1% SDS, 1 mM EDTA pH 8.0, and 10 mM Tris pH 8.0), transferred in a Covaris milliTUBE 1 ml AFA Fiber (Covaris #520135) and sonicated for 12 min using a Covaris ultrasonicator at 5% duty cycle, 140 PIP, and 200 cycles per burst. The sonication cycle was repeated twice. Sonicated chromatin was spun down at 10,000g for 5 min at 4C, the supernatant was collected and moved to a new tube. 20 μl of chromatin was taken to analyze appropriate chromatin shearing on a 1% agarose gel, while 1/10 of the total volume (∼90 μl) was topped up with 5× IP buffer (250 mM HEPES, 1.5 M NaCl, 5 mM EDTA pH 8.0, 5% Triton X-100, 0.5% DOC, and 0.5% SDS) and frozen down at −20°C for total input analysis. The remaining chromatin was topped up to 1ml with 5× IP buffer, then 30 μl of protein A/G Magnetic Beads (Thermo Fischer Scientific #88802) and 3μg of antibody were added to each tube and samples were left to rotate overnight at 4°C.

The following day, beads were washed in 1ml of 1× IP buffer by constant rotation at 4°C for 10 min. This step was repeated twice. Two more washes were performed: the first one in DOC buffer (10 mM Tris pH 8; 0.25 M LiCl; 0.5% NP40; 0.5% DOC; 1 mM EDTA) and the second one in 1x TE buffer. Then, beads were re-suspended in 100 μl of freshly prepared Elution buffer (1% SDS, 0.1M NaHCO3) and agitated constantly on a vortex for 15 min at RT. The eluted chromatin was transferred to a new tube, and elution was repeated again as before by adding 50 μl of Elution buffer to the beads. The eluted chromatin was combined. Finally, 10 μl of 5M NaCl was added to the eluted chromatin as well as to the thawed total input tubes. Samples were incubated overnight at 65°C in a water bath. The next day, the DNA was purified using the Zymo Genomic DNA clean and concentrator kit (Zymo Research #D4011) and eluted in 30 μl of Ultrapure water. For qPCR analysis, samples were handled as described above for CUT&RUN-qPCR. Specifically, a 1:10 dilution was performed and 2 μl of diluted DNA was amplified by means of a QuantStudio 5 (Applied Biosystems) thermal cycler using the SYgreen Blue Mix (PCRbio) and specific primers for both targeted and control genomic regions. Relative abundance of histone marks was determined by using the “percent input” method (the 2^-Ct values obtained from the ChiP samples were divided by the 2^-Ct values of the input samples). Data is then shown as relative fold change between experimental samples and control samples (e.g. CD^scFV^ over GFP^scFV^).

### ATAC-seq

Cells were initially treated in culture medium with 200 U/ml of DNaseI for 30 min at 37°C to digest degraded DNA released from dead cells, and then harvested. Cells were then washed five times in PBS, dissociated with TrpLE and counted. 5 × 10^4^ cells were pelleted at 500g at 4°C for 5 min. The supernatant was removed and cell pellet was resuspended in 50 μl of cold ATAC Resuspension buffer (10 mM Tris-HCl pH7.4, 10 mM NaCl, 3 mM MgCl2, supplemented with 0.1% NP40, 0.1% Tween20 and 0.01% digitonin), followed by incubation on ice for 3 min. Lysis was stopped by washing with 1ml of cold ATAC Resuspension buffer supplemented with 0.1% Tween20 only. Nuclei were pelleted at 500g for 10 min at 4°C. The supernatant was removed and the nuclei were resuspended in 50 μl of transposition mixture (25 μl 2xTD buffer, 2.5 μl transposase from the Illumina Tagment DNA Enzyme and Buffer Kit #20034197, 16.5 μl PBS1x, 0.5 μl 1% digitonin, 0.5 μl 10% tween20 and 5 μl H_2_O). Samples were incubated at at 37°C for 30 min in a thermomixer while shaking at 1,000 RPM. Next, the DNA was purified using the Zymo Genomic DNA clean and concentrator kit (Zymo Research #D4011) and eluted in 21 μl of elution buffer. 20 μl was used for PCR amplification using Q5 hot start high-fidelity polymerase (NEB #M0494S) and a unique combination of the dual-barcoded primers P5 and P7 Nextera XT Index kit (Illumina #15055293). The cycling conditions were: 98°C for 30 s; 98°C for 10 s; 63°C for 30 s; 72°C for 1 min; 72°C for 5 min, repeated for five cycles. After, 5 μl of the pre-amplified mixture was used to determine additional cycles by qPCR amplification using SYgreen Blue Mix (PCRbio) and the P5 and P7 primers selected above in a QuantStudio 5 (Applied Biosystems) thermal cycler. The number of additional PCR cycles to be performed was determined by plotting linear Rn versus cycle and by identifying the cycle number that corresponds to one-third of the maximum fluorescent intensity (Buenrostro et al. 2015). The determined extra PCR cycles were performed by placing the pre-amplified reaction back in the thermal cycler. Finally, clean-up of the amplified library was performed using again the DNA clean and concentration kit (Zymo #D4014) and the DNA was eluted in 20 μl of H_2_O. After quantification and quality check with an automated electrophoresis system (Agilent Tape Station system), library samples were pooled together and sequenced on the Nextseq Illumina sequencing system (paired-end 40 sequencing). Following sequencing, raw reads were first trimmed with TrimGalore (v0.4.3.1, reads > 20 bp and quality > 30) and then quality checked with FastQC (v0.72). The resulting reads were aligned to custom mouse mm10 genome containing the reporter using Bowtie2 (v2.3.4.3, paired-end settings, fragment size 0-1,000, --fr, allow mate dovetailing). Aligned sequences were then analysed with seqmonk (Babraham bioinformatics, v1.47.0) by performing enrichment quantification of the normalised reads.

### Statistical analysis

Details on all statistical analysis used in this paper, including the statistical tests used, the number of replicates and precision measures, are indicated in the corresponding figure legends. Statistical analysis of replicate data was performed using appropriate strategies in Prism GraphPad statistical software (v8.4.3), with the following significance designations: n.s *P>0*.*05, * P ≤ 0*.*05, ** P ≤ 0*.*01, *** P ≤ 0*.*001*.

## Data Accessibility

All data derived from next generation sequencing assays have been deposited in the publically available ArrayExpress database under the accession codes E-MTAB-12103, E-MTAB-12101, E-MTAB-12100.

## ACKNOWLEDGEMENTS

We are grateful to EMBL core facilities, in particular to Gerald Pfister (FCF), Nicolas Descostes (Informatics) and Charles Giradot (GBCS) for experimental and logistics assistance. We thank Lorena Andrade for experimental support. We are also grateful to Mathieu Boulard and Ana Boskovic for critically reading the manuscript. This study was funded by a European Molecular Biology Laboratory (EMBL) programme grant to J.A.H.

## AUTHOUR CONTRIBUTIONS

C.P performed experiments, data analysis, and co-wrote the manuscript. M.M, S.T, and V.C performed key experiments. J.A.H designed and supervised the study, performed data analysis, and wrote the manuscript.

## REFERENCES

1. Kouzarides, T. Chromatin modifications and their function. Cell 128, 693–705 (2007).

2. Blackledge, N.P. & Klose, R.J. The molecular principles of gene regulation by Polycomb repressive complexes. Nat Rev Mol Cell Biol 22, 815–833 (2021).

3. Allshire, R.C. & Madhani, H.D. Ten principles of heterochromatin formation and function. Nat Rev Mol Cell Biol 19, 229–244 (2018).

4. Cavalli, G. & Heard, E. Advances in epigenetics link genetics to the environment and disease. Nature 571, 489–499 (2019).

5. Grosswendt, S. et al. Epigenetic regulator function through mouse gastrulation. Nature 584, 102–108 (2020).

6. Weinberg, D.N. et al. The histone mark H3K36me2 recruits DNMT3A and shapes the intergenic DNA methylation landscape. Nature 573, 281–286 (2019).

7. Barski, A. et al. High-resolution profiling of histone methylations in the human genome. Cell 129, 823–37 (2007).

8. Consortium, E.P. An integrated encyclopedia of DNA elements in the human genome. Nature 489, 57–74 (2012).

9. Roadmap Epigenomics, C. et al. Integrative analysis of 111 reference human epigenomes. Nature 518, 317–30 (2015).

10. Gorkin, D.U. et al. An atlas of dynamic chromatin landscapes in mouse fetal development. Nature 583, 744–751 (2020).

11. Meissner, A. et al. Genome-scale DNA methylation maps of pluripotent and differentiated cells. Nature 454, 766–770 (2008).

12. Dorighi, K.M. et al. Mll3 and Mll4 Facilitate Enhancer RNA Synthesis and Transcription from Promoters Independently of H3K4 Monomethylation. Mol Cell 66, 568–576 e4 (2017).

13. Zhang, T., Zhang, Z., Dong, Q., Xiong, J. & Zhu, B. Histone H3K27 acetylation is dispensable for enhancer activity in mouse embryonic stem cells. Genome Biol 21, 45 (2020).

14. Howe, F.S., Fischl, H., Murray, S.C. & Mellor, J. Is H3K4me3 instructive for transcription activation? Bioessays 39, 1–12 (2017).

15. Wang, Z. et al. Prediction of histone post-translational modification patterns based on nascent transcription data. Nature Genetics 54, 295–305 (2022).

16. O’Carroll, D. et al. The polycomb-group gene Ezh2 is required for early mouse development. Mol Cell Biol 21, 4330–6 (2001).

17. Sankar, A. et al. Histone editing elucidates the functional roles of H3K27 methylation and acetylation in mammals. Nat Genet 54, 754–760 (2022).

18. Liu, H. et al. A method for systematic mapping of protein lysine methylation identifies functions for HP1beta in DNA damage response. Mol Cell 50, 723–35 (2013).

19. Huang, J. et al. p53 is regulated by the lysine demethylase LSD1. Nature 449, 105–8 (2007).

20. Chrysanthou, S. et al. The DNA dioxygenase Tet1 regulates H3K27 modification and embryonic stem cell biology independent of its catalytic activity. Nucleic Acids Res (2022).

21. Jiang, Q. et al. G9a Plays Distinct Roles in Maintaining DNA Methylation, Retrotransposon Silencing, and Chromatin Looping. Cell Rep 33, 108315 (2020).

22. Nakamura, M., Gao, Y., Dominguez, A.A. & Qi, L.S. CRISPR technologies for precise epigenome editing. Nat Cell Biol 23, 11–22 (2021).

23. Policarpi, C., Dabin, J. & Hackett, J.A. Epigenetic editing: Dissecting chromatin function in context. Bioessays 43, e2000316 (2021).

24. Hilton, I.B. et al. Epigenome editing by a CRISPR-Cas9-based acetyltransferase activates genes from promoters and enhancers. Nat Biotechnol 33, 510–7 (2015).

25. Kwon, D.Y., Zhao, Y.T., Lamonica, J.M. & Zhou, Z. Locus-specific histone deacetylation using a synthetic CRISPR-Cas9-based HDAC. Nat Commun 8, 15315 (2017).

26. Cano-Rodriguez, D. et al. Writing of H3K4Me3 overcomes epigenetic silencing in a sustained but context-dependent manner. Nature Communications 7, 12284 (2016).

27. O’Geen, H. et al. dCas9-based epigenome editing suggests acquisition of histone methylation is not sufficient for target gene repression. Nucleic Acids Res 45, 9901–9916 (2017).

28. Kearns, N.A. et al. Functional annotation of native enhancers with a Cas9-histone demethylase fusion. Nat Methods 12, 401–403 (2015).

29. Saunderson, E.A. et al. Hit-and-run epigenetic editing prevents senescence entry in primary breast cells from healthy donors. Nat Commun 8, 1450 (2017).

30. Li, K. et al. Interrogation of enhancer function by enhancer-targeting CRISPR epigenetic editing. Nat Commun 11, 485 (2020).

31. Amabile, A. et al. Inheritable Silencing of Endogenous Genes by Hit-and-Run Targeted Epigenetic Editing. Cell 167, 219–232 e14 (2016).

32. Braun, S.M.G. et al. Rapid and reversible epigenome editing by endogenous chromatin regulators. Nat Commun 8, 560 (2017).

33. Swain, T. et al. A modular dCas9-based recruitment platform for combinatorial epigenome editing. (2022).

34. Zhao, W. et al. Investigating crosstalk between H3K27 acetylation and H3K4 trimethylation in CRISPR/dCas-based epigenome editing and gene activation. Sci Rep 11, 15912 (2021).

35. Millan-Zambrano, G., Burton, A., Bannister, A.J. & Schneider, R. Histone post-translational modifications - cause and consequence of genome function. Nat Rev Genet (2022).

36. Isbel, L., Grand, R.S. & Schübeler, D. Generating specificity in genome regulation through transcription factor sensitivity to chromatin. Nature Reviews Genetics (2022).

37. Morita, S. et al. Targeted DNA demethylation in vivo using dCas9-peptide repeat and scFv-TET1 catalytic domain fusions. Nat Biotechnol 34, 1060–1065 (2016).

38. Carlini, V., Policarpi, C. & Hackett, J.A. Epigenetic inheritance is gated by naive pluripotency and Dppa2. EMBO J, e108677 (2022).

39. Chen, B. et al. Dynamic imaging of genomic loci in living human cells by an optimized CRISPR/Cas system. Cell 155, 1479–91 (2013).

40. Henikoff, S. & Shilatifard, A. Histone modification: cause or cog? Trends Genet 27, 389–96 (2011).

41. Douillet, D. et al. Uncoupling histone H3K4 trimethylation from developmental gene expression via an equilibrium of COMPASS, Polycomb and DNA methylation. Nat Genet 52, 615–625 (2020).

42. Weinert, B.T. et al. Time-Resolved Analysis Reveals Rapid Dynamics and Broad Scope of the CBP/p300 Acetylome. Cell 174, 231–244 e12 (2018).

43. Ong, C.T. & Corces, V.G. CTCF: an architectural protein bridging genome topology and function. Nat Rev Genet 15, 234–46 (2014).

44. Weintraub, A.S. et al. YY1 is a structural regulator of enhancer-promoter loops. Cell 171, 1573–1588. e28 (2017).

45. Lourenco, C. et al. MYC protein interactors in gene transcription and cancer. Nat Rev Cancer 21, 579–591 (2021).

46. Endoh, M. et al. PCGF6-PRC1 suppresses premature differentiation of mouse embryonic stem cells by regulating germ cell-related genes. Elife 6(2017).

47. Lauberth, S.M. et al. H3K4me3 interactions with TAF3 regulate preinitiation complex assembly and selective gene activation. Cell 152, 1021–36 (2013).

48. Santos-Rosa, H. et al. Active genes are tri-methylated at K4 of histone H3. Nature 419, 407–11 (2002).

49. Clouaire, T. et al. Cfp1 integrates both CpG content and gene activity for accurate H3K4me3 deposition in embryonic stem cells. Genes Dev 26, 1714–28 (2012).

50. Margaritis, T. et al. Two distinct repressive mechanisms for histone 3 lysine 4 methylation through promoting 3’-end antisense transcription. PLoS Genet 8, e1002952 (2012).

51. Hu, D. et al. Not All H3K4 Methylations Are Created Equal: Mll2/COMPASS Dependency in Primordial Germ Cell Specification. Mol Cell 65, 460–475 e6 (2017).

52. Vermeulen, M. et al. Selective anchoring of TFIID to nucleosomes by trimethylation of histone H3 lysine 4. Cell 131, 58–69 (2007).

53. Benayoun, B.A. et al. H3K4me3 breadth is linked to cell identity and transcriptional consistency. Cell 158, 673–88 (2014).

54. Domcke, S. et al. Competition between DNA methylation and transcription factors determines binding of NRF1. Nature 528, 575–9 (2015).

55. Grand, R.S. et al. BANP opens chromatin and activates CpG-island-regulated genes. Nature 596, 133–137 (2021).

56. Methot, S.P. et al. H3K9me selectively blocks transcription factor activity and ensures differentiated tissue integrity. Nat Cell Biol 23, 1163–1175 (2021).

57. Halfon, M.S. Perspectives on Gene Regulatory Network Evolution. Trends Genet 33, 436–447 (2017).

58. Do, C. et al. Genetic-epigenetic interactions in cis: a major focus in the post-GWAS era. Genome Biol 18, 120 (2017).

59. Skene, P.J. & Henikoff, S. An efficient targeted nuclease strategy for high-resolution mapping of DNA binding sites. Elife 6(2017).

