## Supplementary Figures for "Systematic Epigenome Editing Captures the Context-dependent Instructive Function of Chromatin Modifications"

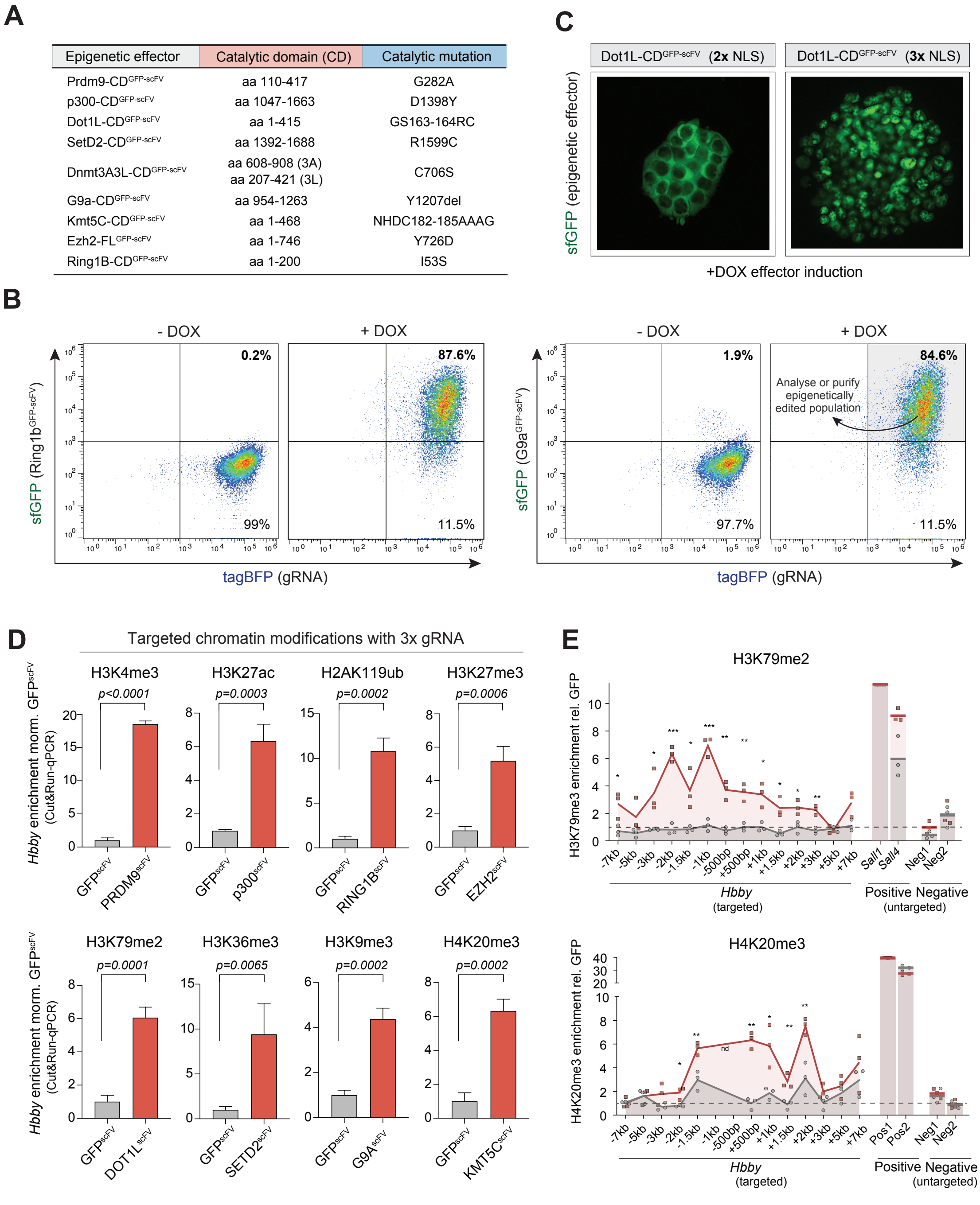

| Epigenetic effector | Catalytic domain (CD) | Catalytic mutation |
|---|---|---|
| Prdm9-CD$^{GFP\text{-}scFV}$ | aa 110-417 | G282A |
| p300-CD$^{GFP\text{-}scFV}$ | aa 1047-1663 | D1398Y |
| Dot1L-CD$^{GFP\text{-}scFV}$ | aa 1-415 | GS163-164RC |
| SetD2-CD$^{GFP\text{-}scFV}$ | aa 1392-1688 | R1599C |
| Dnmt3A3L-CD$^{GFP\text{-}scFV}$ | aa 608-908 (3A) aa 207-421 (3L) | C706S |
| G9a-CD$^{GFP\text{-}scFV}$ | aa 954-1263 | Y1207del |
| Kmt5C-CD$^{GFP\text{-}scFV}$ | aa 1-468 | NHDC182-185AAAG |
| Ezh2-FL$^{GFP\text{-}scFV}$ | aa 1-746 | Y726D |
| Ring1B-CD$^{GFP\text{-}scFV}$ | aa 1-200 | I53S |



**Supplementary Figure 1. An optimised toolkit for dynamic & precision chromatin state perturbations.**
(**A**) Table detailing the catalytic domains (CD) used as epigenetic 'effectors' in this study, and the precise point-mutant controls to specifically disrupt their catalytic activity. Each CD effector is tagged with superfolder GFP (sfGFP) and an scFV domain that specifically binds the GCN4 tail of dCas9$^{GCN4}$. (**B**) Representative flow cytometry dot plots showing DOX-dependent induction of the epigenetic editing systems that programme H2AK119ub (left) and H3K9me2 (right). The enhanced gRNA is constitutively expressed and marked by tagBFP (x-axis). dCas9$^{GCN4}$ and each CD$^{scFV}$ effector is activated by DOX, leading to nuclear GFP signal and epigenetic editing. Note GFP signal confirms CD$^{scFV}$ or mut-CD$^{scFV}$ stability, enables dose-dependent responses to be ascertained, and is used to flow sort pure populations of cells that have appropriately activated the editing system (GFP+). (**C**) Representative image showing that CD$^{scFV}$ effectors often required additional nuclear localization sequences (NLS) for nuclear accumulation and efficient epigenetic editing. (**D**) Bar plot showing enrichment of the indicated chromatin modifications at the endogenous *Hbby* locus, following targeted editing with the relevant CD$^{scFV}$ or control GFP$^{scFV}$ targeting. These data were obtained with three concomitant gRNA and independently of the data with a single gRNA shown in Figure 1B. (**E**) High resolution enrichment of H3K79me2 (upper) and H4K20me3 (lower) across the entire *Hbby* locus targeted with three gRNAs. Enrichment at positive control endogenous loci and negative control (untargeted) loci is shown.

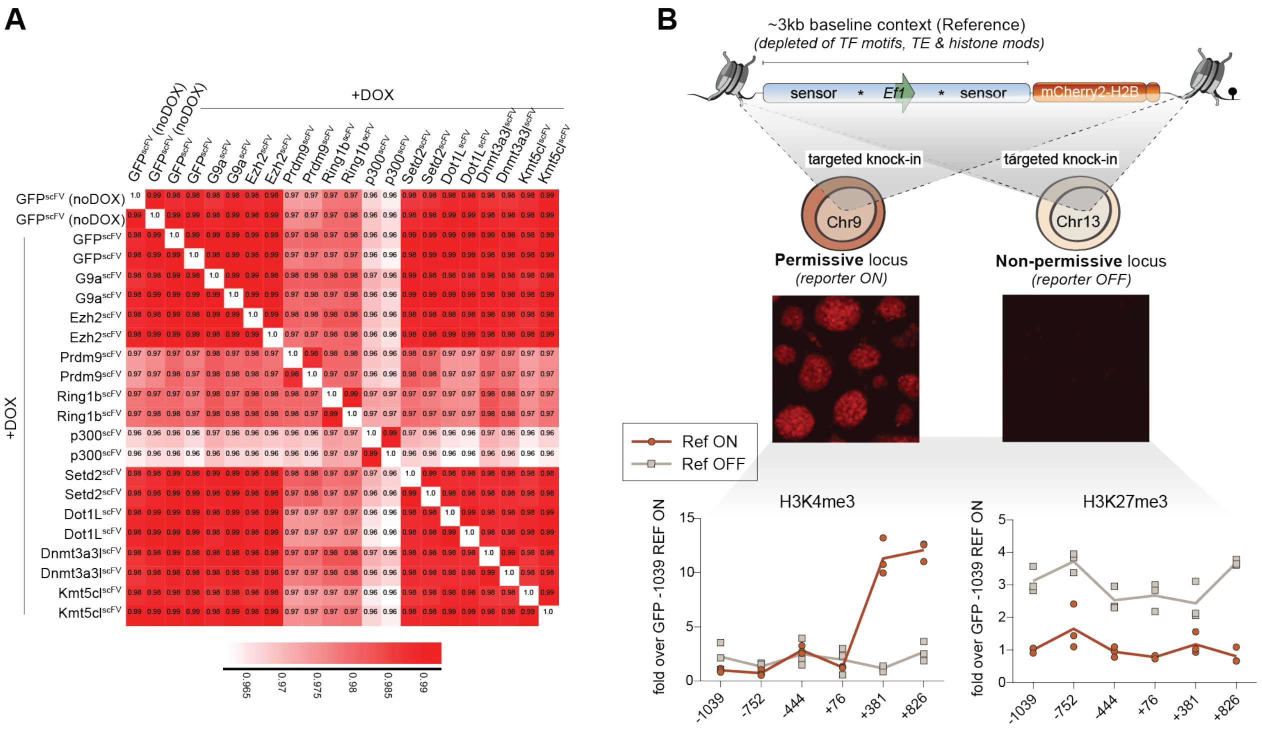


**Supplementary Figure 2. Reporter (epi)genomic contexts & minimal OFF-targeting.**
(**A**) Correlation matrix of replicate whole transcriptomes following induction of the indicated epigenetic editing system with DOX. We routinely observed high correlation (>0.98) between sample global expression, with minimal OFF-target mis-expression, indicative of high ON-target activity. The exception is p300$^{scFV}$, and we therefore reduced the DOX concentration to mitigate indirect effects. (**B**) Quantification of acquired H3K4me3 and H3K27me3 at identical reference reporters located within two distinct genomic contexts (see schematic above).

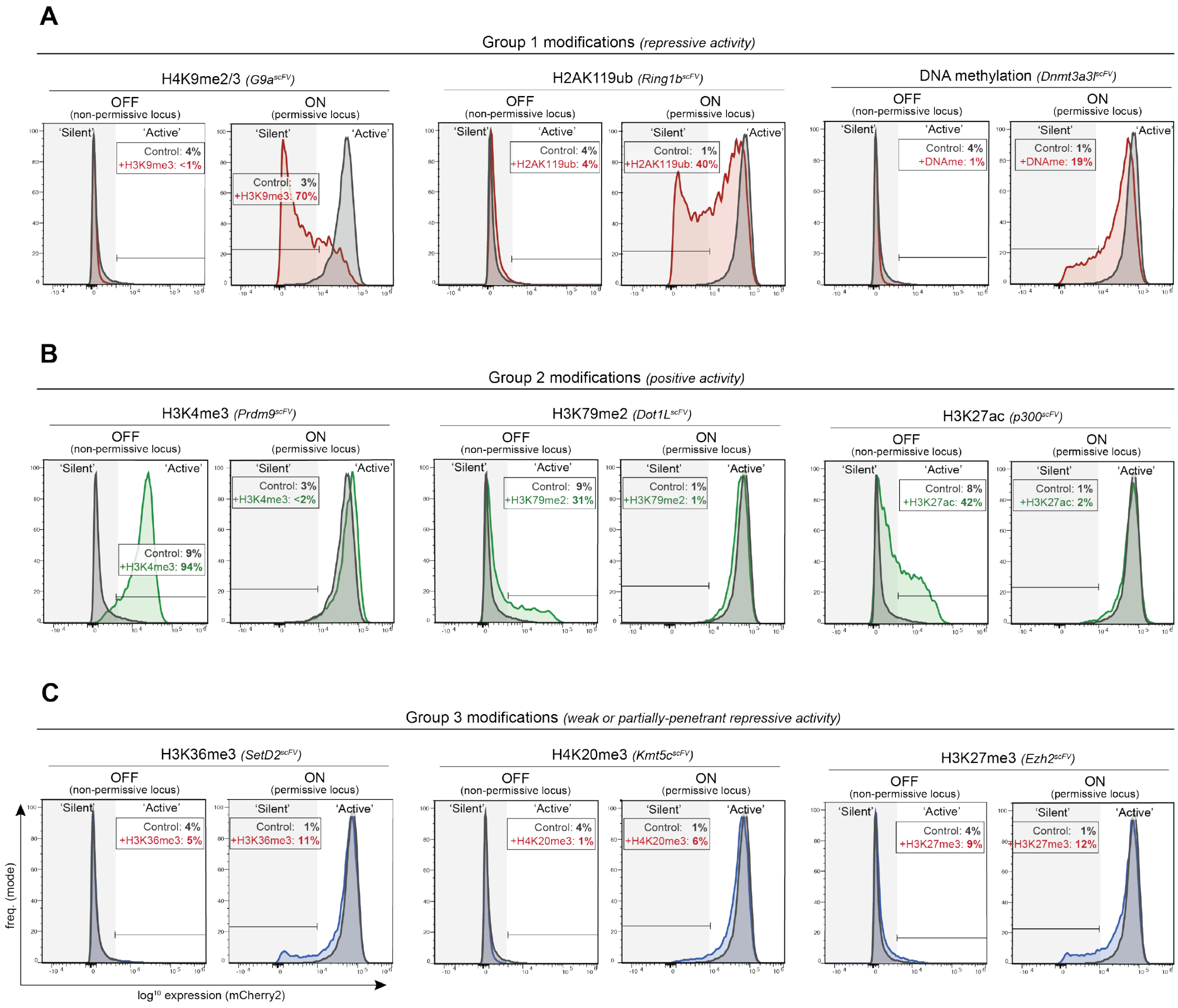


**Supplementary Figure 3. Impact of programmed chromatin modifications on distribution of gene expression from distinct loci.**
(**A-C**) Representative flow cytometry histograms of reporter gene expression following *de novo* programming of the indicated chromatin modification. For each modification, the transcriptional effect is shown from a non-permissive location (initial expression OFF; see left panels) and on an identical promoter in a permissive location (initial expression ON; see right panels). The percentage of cells that acquire a new expression state following precision chromatin editing in each context is indicated, along with control (GFP$^{scFV}$) targeting. Based on reproducible transcriptional responses, we grouped chromatin modifications into functional cohorts whereby (**A**) deposition promotes significant gene repression amongst a major fraction of cells, from an active genomic location (**B**) deposition facilitates significant gene activation amongst a major fraction of cells, from a repressed location, and (**C**) *de novo* targeting has a subtle or highly partially-penetrant repressive effect.

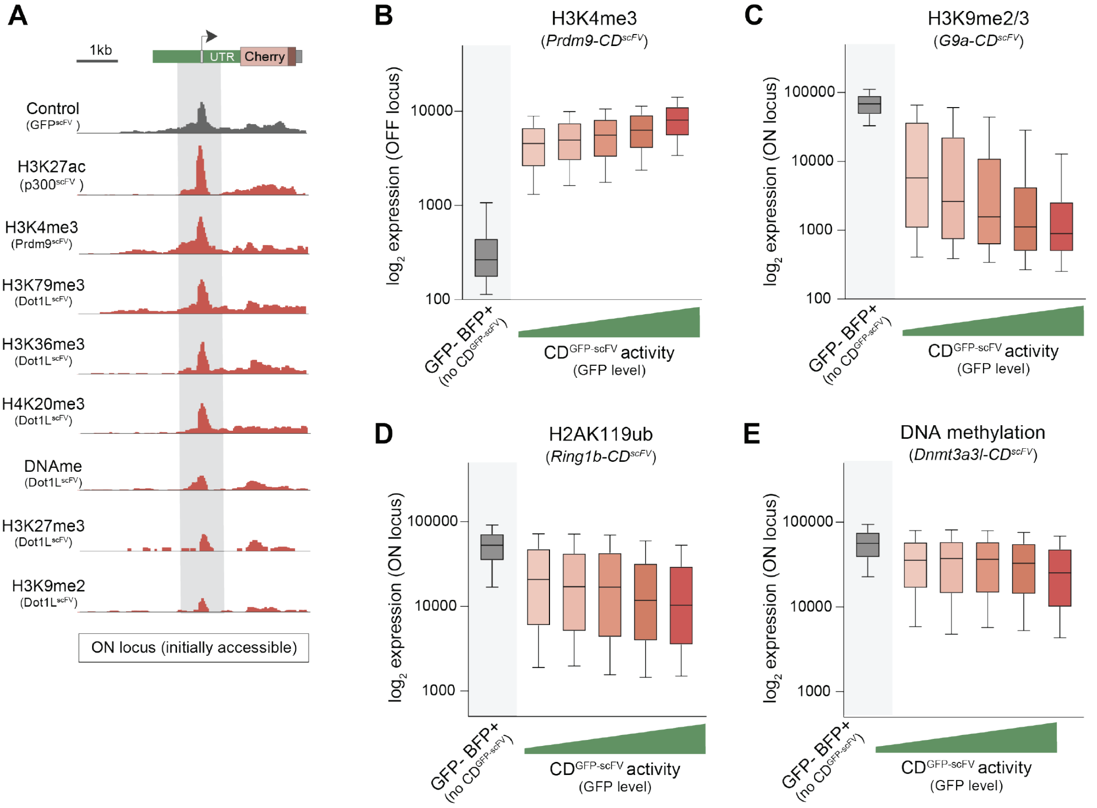


**Supplementary Figure 4. Dose-dependent impact of programmed histone modifications.**
(**A**) Promoter accessibility at the permissive reporter locus measured by ATAC-seq. Shown is the genome view of promoter accessibility following *de novo* programming of the indicated chromatin modification. (**B-E**) Dose-dependent transcriptional responses to the indicated chromatin modification effectors. A single population of +DOX cells was stratified based on the level of induced $CD^{scFV}$ expression, as determined by GFP . Shown is the transcriptional response of the reporter, which is directly correlated with the amount of epigenetic editing activity in the cell. Representative dose-dependent responses are displayed as boxplots of single-cell expression levels following programming of **(B)** H3K4me3 **(C)** H3K9me2/3 **(D)** H2AK119ub **(E)** DNA methylation.

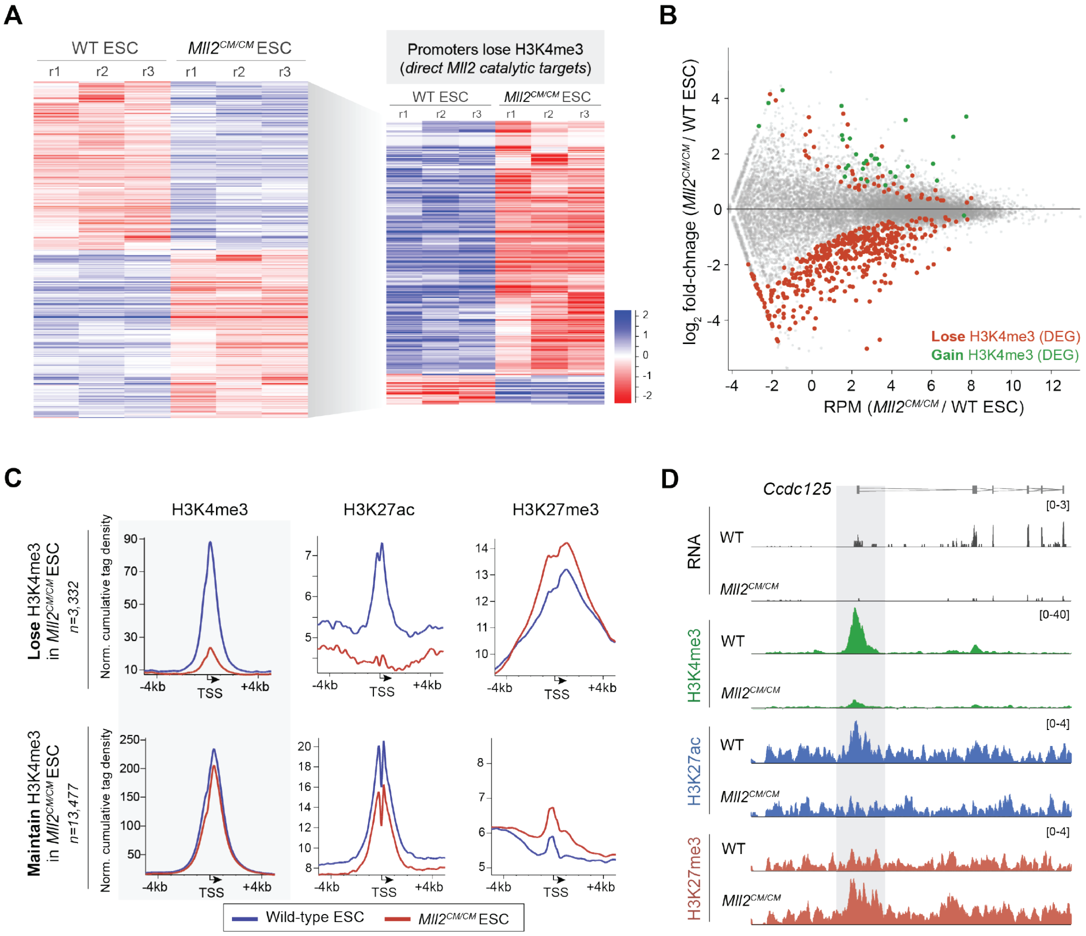


**Supplementary Figure 5. Analysis of Mll2$^{CM/CM}$ ESC that specifically lack H3K4me3 methylase activity.**

(**A**) Left: Heatmap showing all differentially expressed genes (DEG; P(adj)<0.05) in *Mll2$^{CM/CM}$* ESC. Right: Heatmap of all DEGs that also become significantly depleted of promoter H3K4me3 in *Mll2$^{CM/CM}$* ESC, thereby representing direct MLL2 targets rather than indirect secondary changes. Note 90% of direct targets are downregulated upon H3K4me3 loss. (**B**) MA plot showing all DEGs that lose H3K4me3 and all DEGs that gain H3K4me3. (**C**) Metaplot of enrichment of the indicated modification over promoters. Shown are all promoters that lose H3K4me3 (upper) and all those that retain H3K4me3 (lower) in *Mll2$^{CM/CM}$* ESC. As a consequence of specific loss of H3K4me3 there is a concomitant loss of H3K27ac and gain in H3K27me3. (**D**) Representative genome view showing expression (RNA), and changes in chromatin marks H3K4me3, H3K27ac, and H3K27me3 upon specific loss of H3K4me3 in *Mll2$^{CM/CM}$* ESC.

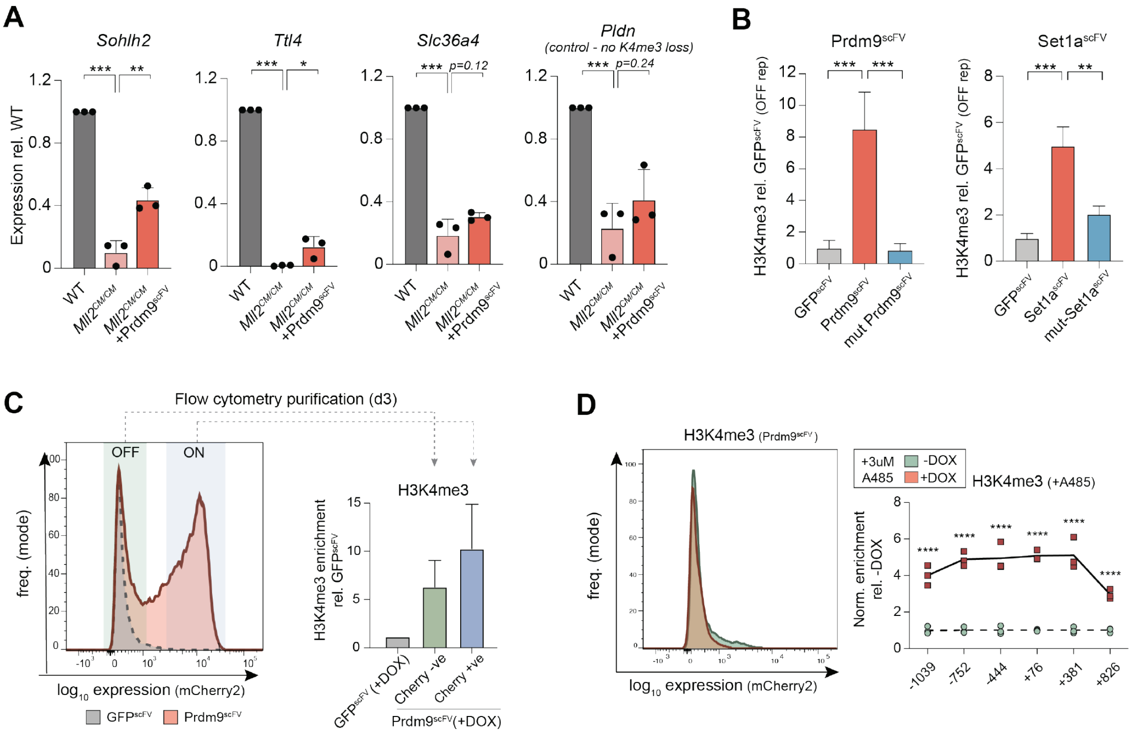


**Supplementary Figure 6. Programming H3K4me3 activates gene expression via H3K27ac.**

(**A**) Re-targeting H3K4me3 back to endogenous promoters that have lost H3K4me3 in *Mll2*$^{CM/CM}$ ESC partially rescues their expression level (see also Fig3B). The control *Pldn* gene exhibit no initial loss of H3K4me3 (indirectly affected), and accordingly was not rescued by deposition of further H3K4me3. (**B**) CUT&RUN-qPCR showing the level of H3K4me3 deposited at the OFF reporter promoter by each of two independent K4me3 effectors and their respective catalytic mutant controls. (**C**) Flow cytometry plot at day 3 of Prdm9$^{scFV}$ induction, showing ~half the population have initiated a transcriptional response (activation). Active (ON) and inactive (OFF) populations were purified and the level of deposited H3K4me3 assayed by CUT&RUN-qPCR. Whilst all cells are enriched with H3K4me3, those with the higher level are active, indicating a threshold level of H3K4me3 is necessary to trigger transcriptional activation. (**D**) Representative flow cytometry histogram showing that programming H3K4me3 no longer activates expression in the presence of an acetylation inhibitor (A485) - compare with S6C at the equivalent timepoint. Shown right is CUT&RUN-qPCR revealing H3K4me3 is programmed in the presence of A485 but cannot elicit downstream effects on transcription.

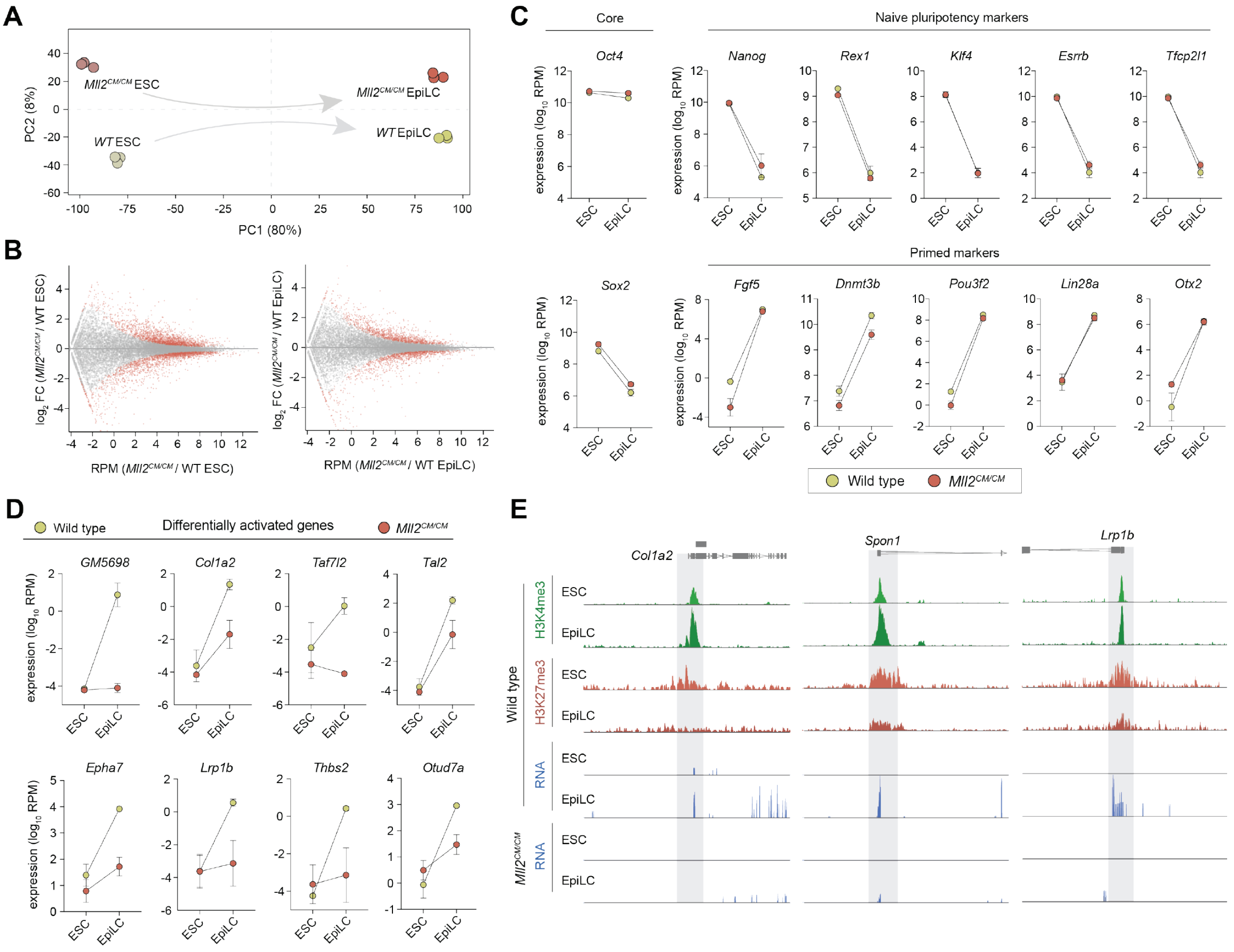


**Supplementary Figure 7. Role of H3K4me3 in *de novo* gene activation during cell fate transition.**
(**A**) Principal component analysis (PCA) of all expressed genes (RPM>1) in WT and *Mll2*$^{CM/CM}$ ESC, and upon transition to EpiLC. (**B**) MA plot showing all DEG in ESC and EpiLC that specifically lack H3K4me3. (**C**) Expression of representative marker genes showing removal of H3K4me3 by *Mll2*$^{CM/CM}$ does not impact the expression of pluripotency and primed (early differentiation) genes. This indicates that *Mll2*$^{CM/CM}$ ESC are fully competent to generate EpiLC, and that any expression changes are not indicative of impaired cell fate commitment. (**D**) Expression of representative differentially expressed genes that fail to activate without H3K4me3 in *Mll2*$^{CM/CM}$ EpiLC. (**E**) Genome view plots showing genes that are activated in WT EpiLC fail to initiate expression in *Mll2*$^{CM/CM}$ EpiLC. These genes normally gain H3K4me3 and lose H3K27me in EpiLC.

**A**

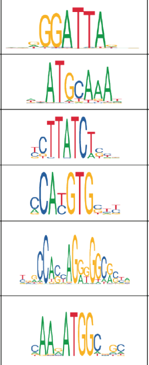

| Reporter | PWM | Inserted motifs |
|---|---|---|
| OTX2 | | 5'-GGGATTAAT-3' |
| OCT4 | | 5'-ATGCAAAT-3' |
| GATA4 | | 5'-TCTTATCTCC-3' |
| MYC | | 5'-CCACGTGCTT-3' |
| CTCF | | 5'-TGGCCAGCAGGGGGCGC-3'<br>5'-CCAGTAGGGGGCGC-3'<br>5'-CCACCAGGGGGCGCTA-3'<br>5'-CCACTAGGGGGCGC-3' |
| YY1 | | 5'-CAAGATGGCTGC-3'<br>5'-CAAGATGGCGGC-3'<br>5'-CAAGATGGCTGC-3'<br>5'-CCAGATGGCCGC-3' |
| G4-U | from *Vlasenok et al.* 2018 | 5'-GGGGATACTGGGGTTC ATGGGGATCCAGGGG-3'<br>5'-GGGGACAGGGGTATGG GGAGGGG-3' |
| G4-D | from *Vlasenok et al.* 2018 | 5'-GGGGATACTGGGGTTC ATGGGGATCCAGGGG-3'<br>5'-GGGGACAGGGGTATGG GGAGGGG-3' |

**B**

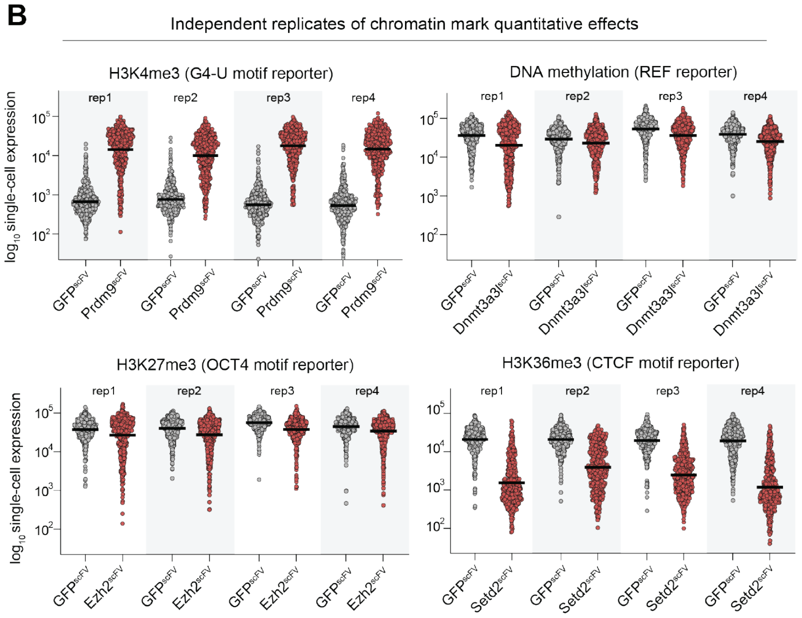


**C**

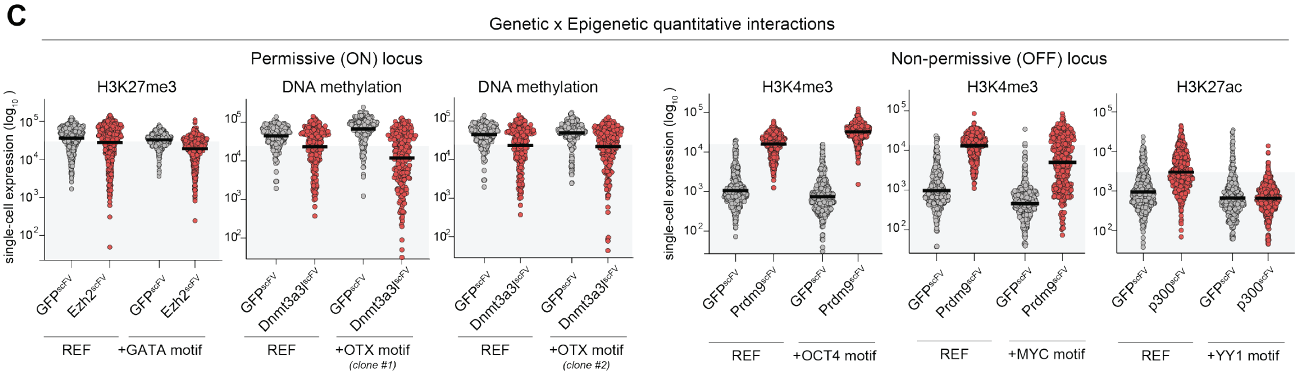


**SuSupplementary Figure 8. Reproducible genetic x epigenetic interactions.**
(**A**) Table illustrating the position weight matrix for each motif deployed across the reporter series, along with the actual inserted motif(s). (**B**) Dot plots showing $log_{10}$ expression in single cells. Displayed within each plot are four independent replicates of programming the same specific epigenetic modification to the same specific reporter, relative to control $GFP^{scFV}$. Different marks and reporter combinations are selected to illustrate the reproducibility of both subtle and major quantitative effects elicited by epigenetic editing. (**C**) Examples of functional interplay between the quantitative impact of a programmed modification and the presence of an underlying TF motif in the reporter. For example, H3K27ac-mediated activation is attenuated in the context of YY1 motifs, H3K4me3 activation is strengthened in the presence of OCT4 motifs, whilst OTX motifs may enhance DNA methylation mediated repression, albeit this effect was not reproducible across all independent clones (representative samples shown).

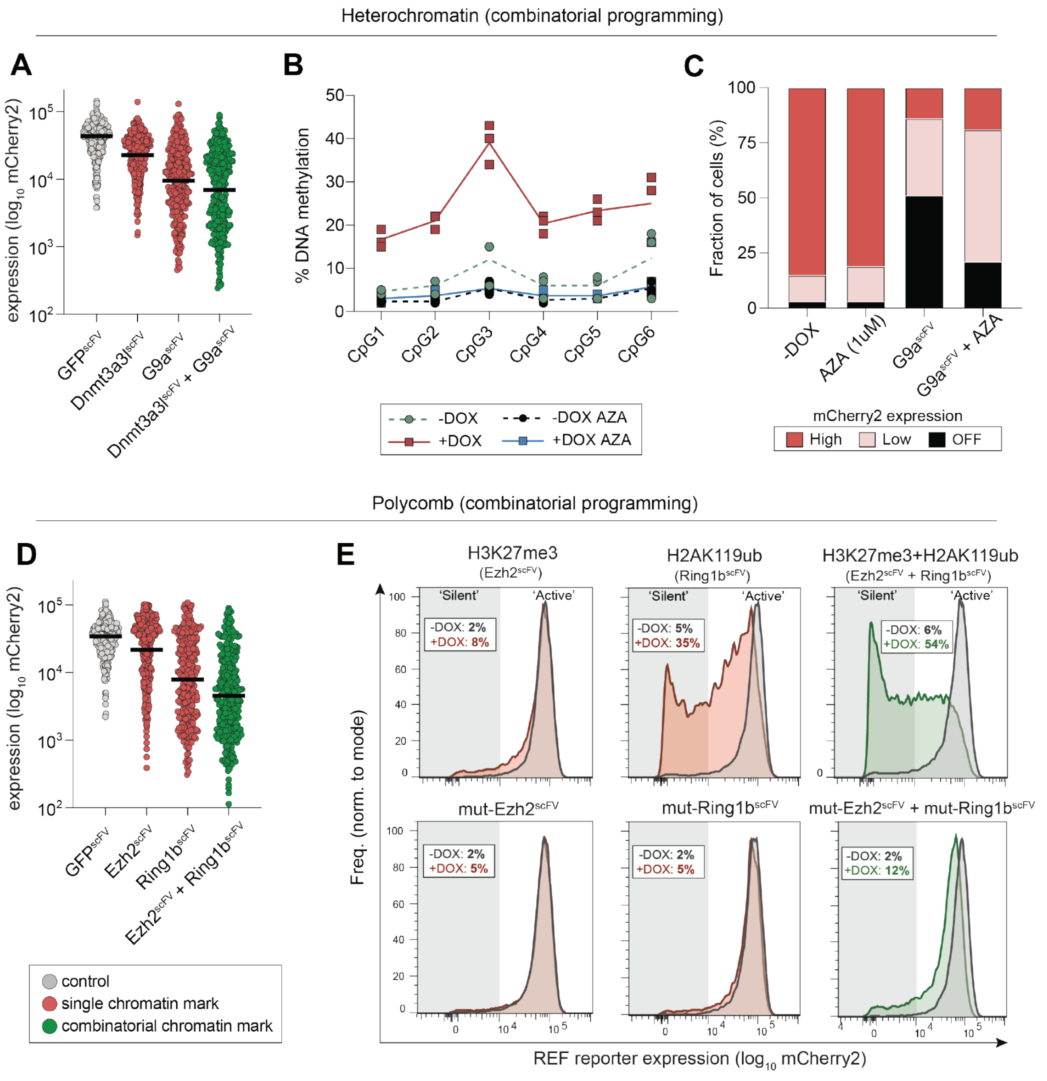


**Supplementary Figure 9. Combinatorial chromatin modifications enhance robustness of single-cell silencing.**
(**A**) Dot plot showing $log_{10}$ single-cell expression upon specific programming of DNA methylation, H3K9me2/3 or both modifications together relative to control. (**B**) DNA methylation pyrosequencing confirming that treatment with 5-azacytdine (1μM) (AZA) impairs DNA methylation deposition at the reporter. (**C**) Fraction of cellular population that is in either a 'off', 'low' or 'high' expression state following epigenetic editing +/- AZA. (**D**) Dot plot showing $log_{10}$ single-cell expression upon specific programming of H3K27me3, H2AK119ub or both polycomb modifications together relative to control. (**E**) Representative and independent flow cytometry plots showing the distribution of gene expression across the population upon single- or combinatorial- polycomb targeting (upper), or with catalytic mutant controls (below). Note that both polycomb marks together increase the penetrance of 'full' silencing.
